# *C. elegans* episodic swimming is driven by multifractal kinetics

**DOI:** 10.1101/2020.04.22.056606

**Authors:** Yusaku Ikeda, Peter Jurica, Hiroshi Kimura, Hiroaki Takagi, Struzik Zbigniew, Ken Kiyono, Yukinobu Arata, Yasushi Sako

## Abstract

Fractal scaling is a common property of temporal change in various modes of animal behavior. The molecular mechanisms of fractal scaling in animal behaviors remain largely unexplored. The nematode *C. elegans* alternates between swimming and resting states in a liquid solution. Here, we report that *C. elegans* episodic swimming is characterized by scale-free kinetics with long-range temporal correlation and local temporal clusterization, which is characterized as multifractal kinetics. Residence times in actively-moving and inactive states were distributed in a power law-based scale-free manner. Multifractal analysis showed that temporal correlation and temporal clusterization were distinct between the actively-moving state and the inactive state. These results indicate that *C. elegans* episodic swimming is driven by transition between two behavioral states, in which each of two transition kinetics follows distinct multifractal kinetics. We found that a conserved behavioral modulator, cyclic GMP dependent kinase (PKG) may regulate the multifractal kinetics underlying an animal behavior. Our combinatorial analysis approach involving molecular genetics and kinetics provides a platform for the molecular dissection of the fractal nature of physiological and behavioral phenomena.

## Introduction

Animal behaviors are organized over a broad range of time scales, ranging from seconds to years, including expansive timescales over lifespan phases, such as infant, juvenile, adult, and elderly phases. Among the temporally organized animal behaviors are rhythmic behaviors characterized by their frequencies. Molecular regulators e.g. for daily rhythmic and oscillatory changes of animal behaviors i.e. for circadian rhythm have been identified and are shown to be regulated by a feedback control^2,3^ Contrastingly, many arrhythmic changes of behavioral and physiological activities in a great variety of animal species, including humans are reported to show self-similar and scale-free structures, which is an indicative for fractal scaling^4–6^. Fractal geometry provides a mathematical framework for characterizing scale-free and self-similar patterns^4–6^. Fractal temporal patterns of behavior and physiology have been reported in the locomotion of birds, mosquito larvae, and flies^7–9^, crawling of cultured *C. elegans* worms^10^, clicking sounds produced by feeding sea horses^11^, and swimming of zooplankton^12,13^. In humans, temporal fractal patterns have been observed in wrist movements during habitual sleep/wake cycles^14^, gait^15,16^, heartbeats^17,18^, and brain activity^19,20^. Altered fractal patterns of activity have been associated with pathological conditions and aging^14–16,18,21,22^, indicating that the fractal patterning in biological activity is a critical measure for characterizing physiological status. Thus, fractal scaling is a clue to study molecular and physiological basis of arrhythmic and complex animal behaviors, and there may be molecular and physiological mechanisms that adhere to the basic principles of fractal geometry. However, the molecular and physiological mechanisms remain unknown.

Owing to its simple anatomy and the availability of a range of genetic tools, *C. elegans* is a powerful model organism for the study of the molecular bases of behavior. In solution, *C. elegans* worms alternate between swimming and resting states on a minute to hour time scale^23,24^. In the swimming state, they alternate between continuously beating their bodies and resting with or without food^23–25^. In the resting state, they maintain a characteristic sharply-bent posture^23^. Episodic swimming is conserved in nematodes cultured in a liquid solution^23^. On a solid agar plate, *C. elegans* also move in an episodic manner, wherein they crawl actively and persistently in one direction or crawl slowly and stay within a small area, behavioral states called roaming and dwelling/quiescence, respectively^24,26,27^ Individual *Drosophila* flies^9^ and *Leptothorax allardycei* worker ants^28^ also alternate between an actively-moving state and a resting state in an episodic manner. Thus, episodic behavior is a conserved presentation in invertebrates that is thought to be adaptive for supporting food exploration, energy conservation, and reproductive success^9,23,24,29^.

In *C. elegans,* episodic swimming is regulated by *egl-4,* which encodes cGMP-dependent kinase (PKG). PKG is a behavior modulating enzyme conserved across invertebrates and vertebrates^30,31^. *C. elegans egl-4/pkg* mutants roam continuously on a solid agar plate and swim when in a solution with less frequent resting than is exhibited by wild-type animals^23,25–27^. Because *egl-4/pkg-dependent* regulation is found in both medium conditions^23,25–27^, *C. elegans* episodic motions in both conditions are thought to be regulated by the same molecular/physiological mechanism^24^. In *Drosophila*, the *foraging/pkg* homolog of *egl-4* had been discovered from natural behavioral polymorphisms wherein flies tend to travel long distances (*rover*) or be sedentary (*sitter*)^32,33^. The expression level of *foraging/pkg* differs between *rover* flies and *sitter* flies and *Drosophila* traveling behavior can be switched by genetic manipulation of *foraging/pkg*^34^ Foraging behaviors in social insects— including those exhibited by honey bees (*Apis mellifera*) and ants (*Pheidole pallidula*) that are determined by developmental stage^35^ and social caste^36^, respectively—are also associated with PKG expression and activity. Thus, PKG is considered to be a conserved modulator of animal behavior^30,31^.

To study scaling of behavior across a broad range of time scales, it is necessary to obtain behavioral activity time series of individual animals for an extended period of time at a high temporal resolution. In this study, we recorded the swimming behavior of 108 individual *C. elegans* at a semi-video rate for a week-long period by individually culturing the worms in a newly developed microfluidic device. The obtained time series data encompassing 10^7^ time points was submitted to multifractal analysis. We found that *C. elegans* episodic swimming is a scale-free process and the transition between actively-moving and inactive states is driven in multifractal kinetic. Additionally, we examined the sensitivity of the multifractal kinetics of the behavior to modulation by PKG. A discussion of potential molecular and physiological mechanisms underlying multifractality in behavior is provided on the basis of previously reported mathematical and physical models.

## Results

### *C. elegans* episodic swimming is a multi-time scale process

Swimming of multiple adult *C. elegans* individuals was monitored in a newly developed microfluidic device, called WormFlo, which is equipped with an array of 108 disc-shaped chambers for culturing individual *C. elegans* (Fig. 1A and B). *C. elegans* individuals were maintained alone in chambers under controlled chemical, temperature, and light intensity conditions (Fig. S2, Fig. S3, and Methods). We recorded swimming at 10^7^ time points with a 50-ms interval (138 h ≈ 5.8 d), and quantified swimming activity with a pixel counting method (Fig. 2A, Fig. S4, Movie S1, Methods, and Supplemental Information). In the absence of an energy source, the animals’ swimming activity diminished gradually over time (Fig. 2B and Movie S2). The swimming activity in WormFlo chambers decayed with kinetics similar to that seen in animals cultured without an energy source in substantially larger (2× diameter, 23 × volume) 96-well plate wells (Movie S3); and the activity was sustained in animals cultured in WormFlo with an energy source (glucose and cholesterol) (Fig. S5A and Movie S4). Therefore, the activity decay observed can be attributed primarily to a physiological response to long-term cultivation without an energy source rather than to physical damage or spatial restriction.

**Figure 1.**
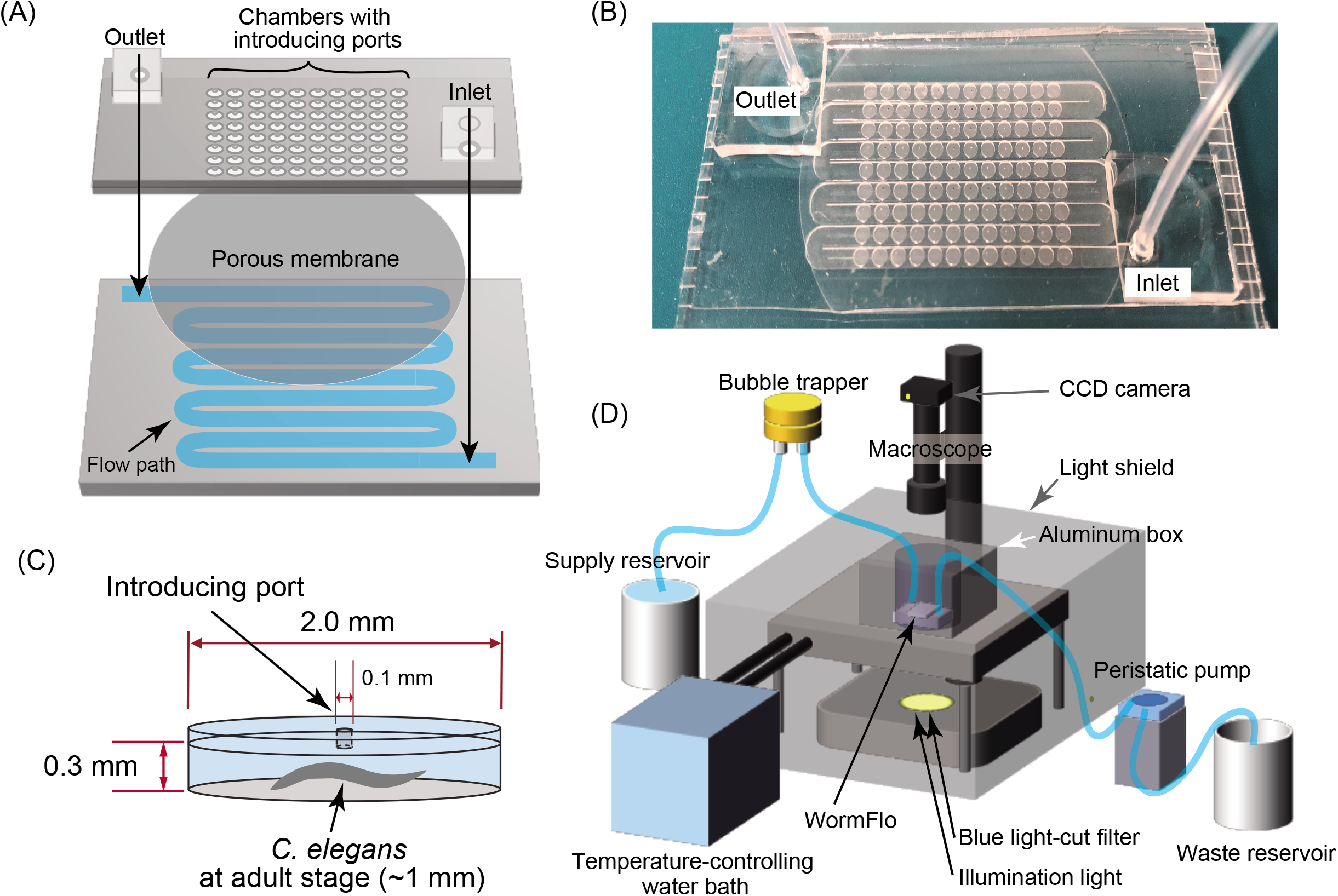
Culturing and recording system for individual *C. elegans* animals. (A and B) The WormFlo apparatus has a vertically two-compartment structure, wherein an array of 108 culture chambers and loading ports of the upper PDMS chip are partitioned from the buffer flow path located in the lower PDMS chip by a porous membrane (details in the Methods). M9 buffer was supplied from the inlet and withdrawn from the outlet. (C) The diameter and height of each culture chamber in the upper PDMS chip were 2 mm and 0.3 mm, approximately 2-fold longer and 3-fold thicker than the ~1-mm-long and ~ 0.1-mm wide *C. elegans* body, respectively. Animals are introduced in each chamber via a 0.1-mm-wide loading port on the roof of a discshaped culture chamber, which was closed with a thin PDMS sheet before M9 buffer was perfused. (D) *C. elegans* animals cultured in the WormFlo are monitored by a macroscope with a CCD camera. Buffer perfusion is driven by a peristatic pump. To avoid loss of water due to evaporation, the WormFlo was submerged in a 15-cm-diameter glass dish (depth, 5.4 cm) filled with M9 buffer. The temperature of the M9-filled 15-cm glass dish was maintained by circulating temperature-controlled water in a water flow path in the attached aluminum block under the 15-cm glass dish (Fig. S3A). Animals cultured in the microfluidic device are illuminated by blue-light filtered light from a halogen lamp in a light shielding box that buffers against light intensity changes due to daily lab activity.

Comparing the activity decay kinetics over the 6-d observation period among individuals led us to define three empirical activity classes: High, long-term high activity; Middle, intermediate-term high activity; and Low, short-term high activity (Fig. 2B). The early and late culturing periods were defined as Pre-starved and Starved time regimes, each of which has high and low swimming activities (for quantitative criteria, see Fig. 2B legend, and Methods). Consistent with previous studies, we observed episodic swimming bouts on minute to hour time scales (Fig. 2C; 8-min scale and 80-min scale), and a characteristic kinked posture during the resting state in the Pre-starved regime (data not shown)^23,24^.

**Figure 2.**
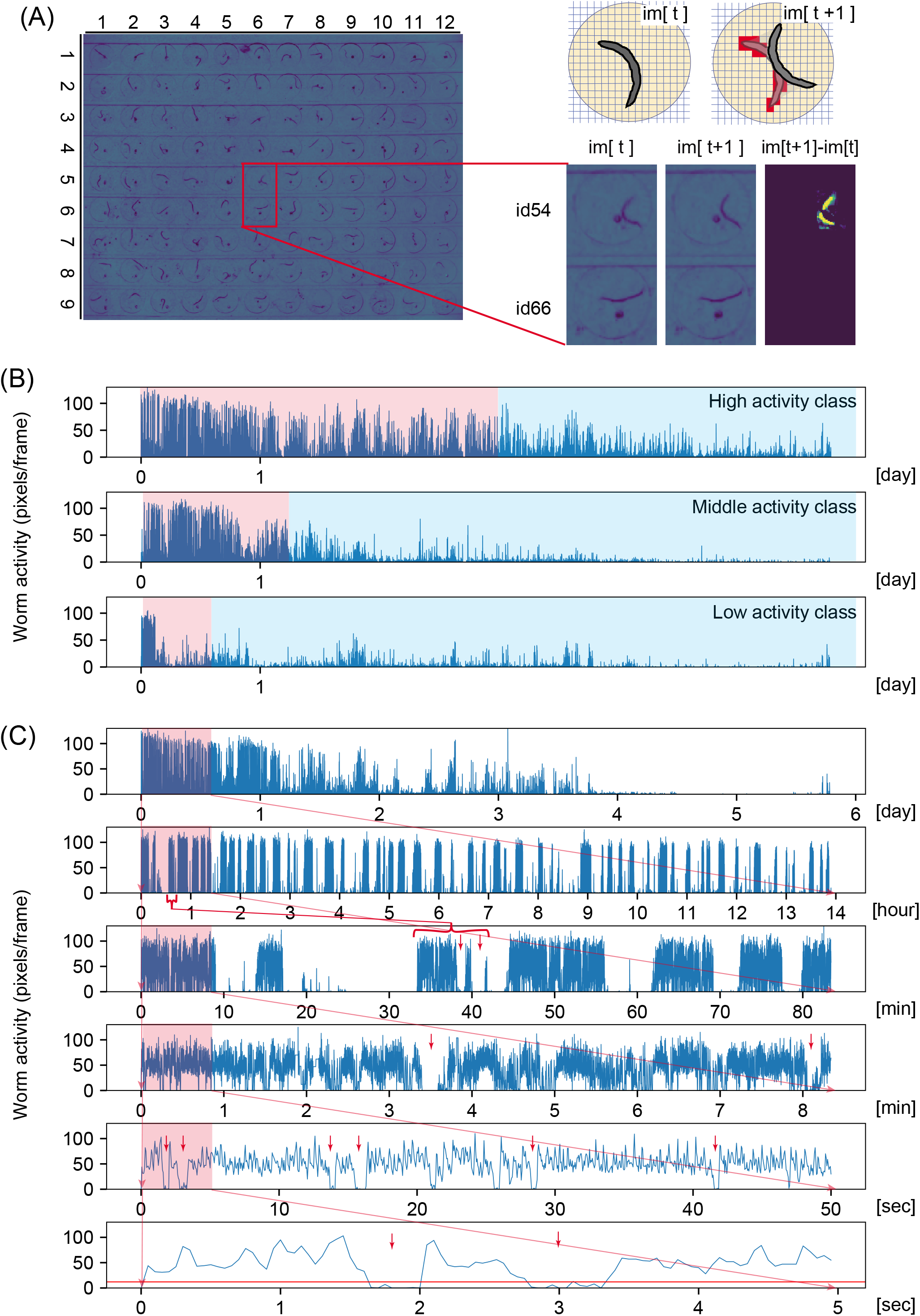
*C. elegans* episodic swimming exhibited a multi-time scale kinetics. Swimming activity of animals cultured in individual chambers was quantified by a pixel counting method (Methods). (A) Chambers in WormFlo are shown with row-column indexes (left figure). Active pixels (intensity difference > 12 in the range of −256 to 256) between time point [t] and time point [t+1] are shown in red pixels in the upper right image and in yellow pixels in the lower right image, respectively. The animal in chamber id 54 moved while the animal in culture chamber id 66 did not move from time [t] to [t+1]. (B) Active pixel numbers are shown on the y-axis as an index of swimming activity with culture time is shown on the x-axis. Six-day temporal activity decay patterns were classified into High, Middle, and Low activity classes by two criteria; average activities during the early half of the recording period (first 3 days) and the ratio of average activity in the early half to that in the late half of the recording period (see Methods). Pre-starved regime (red area) and Starved regime (blue area) were defined for each activity class. For the High activity class, the Pre-starved and Starved regimes were defined by early 50% period from the start of recording through the early half of the recording period and the remainder of the period. For the Middle activity class, Pre-starved regime and Starved regime were defined by the period from the start of recording to early 20% period of the recording period and the remainder. For the Low activity class, the Pre-starved and Starved regimes were defined by the period from the start of recording to early 10% of the recording period and the remainder. (C) Swimming activities of a representative animal in various time scales; full recording time scale (10^7^ timepoints) at the top. Each of the lower panels are a 10× magnification of the first tenth of its upper panel (red area). Activity threshold at 12 pixels/frame is shown by a red horizontal line in the 5-s scale. Animal activities at 6 d-, 1 d-, 1 h-, 10 min-, 1 min-, and 1 s-scales are shown.

Interestingly, we observed a swimming bout cluster on a 1 day-scale that could be divided in several clusters in a magnified view of about 1 h (Fig. 2C, compare the red bracketed region in the 14-h scale vs. red arrows in the 80-min scale). This nested temporal structure was observed repeatedly over a series of magnifications from a 1-h scale to a 10-min scale (Fig. 2C, red region in 80-min scale vs. red arrows in 8-min scale), and between the 1-min and 1-s scales (Fig. 2C, red bracket region in 8-min scale vs. red arrows in 50-s and 5-s scales). In the 1-s scale, animals alternated between bending their bodies and beating their bodies for swimming (Movie S1). Series of beating motions were interrupted with intermittent short resting periods, referred as to “posing” (Movie S1). During posing, they were transiently motionless in a bent posture (Movie S1, see legend). The temporal resolution at which continuous shape could be detected in H265 codec-compressed movies was 0.25 s (Fig. S6 and Supplemental Information), which was sufficient to detect subsecond “posing” on a 1-s scale. The temporally nested structure of *C. elegans* episodic motion was observed on time scales of a magnification range of about 1,000 times. Thus, *C. elegans* episodic swimming has multi-timescale dynamics with a self-similar temporal structure.

### A scale-free property in *C. elegans* episodic swimming

Continuous swimming ceased and resumed suddenly (Fig. 2C). Consistently, we observed a bimodal probability distribution of activity strength (Fig. 3A), indicating that *C. elegans* episodic swimming can be characterized by a two-state transition model between actively-moving and inactive states. To reveal the kinetics of the state transition, we studied the statistical distribution of residence times in active and inactive states. Active and inactive states were defined as periods above or below an activity threshold, which was the value in the bimodal distribution valley (Fig. 3A) of swimming activity time series (red horizontal line in Fig. 2C, 5-s scale). Residence time in the two states were obtained alternately from swimming activity time series; eventually, we obtained round series data for active and inactive states (Fig. 3B and C). Residence times in the two states were distributed along a broad temporal range, ranging from subsecond to 10-s orders for the active state and ranging from subsecond to 100-s orders for the inactive state (Fig. 3B and C).

**Figure 3.**
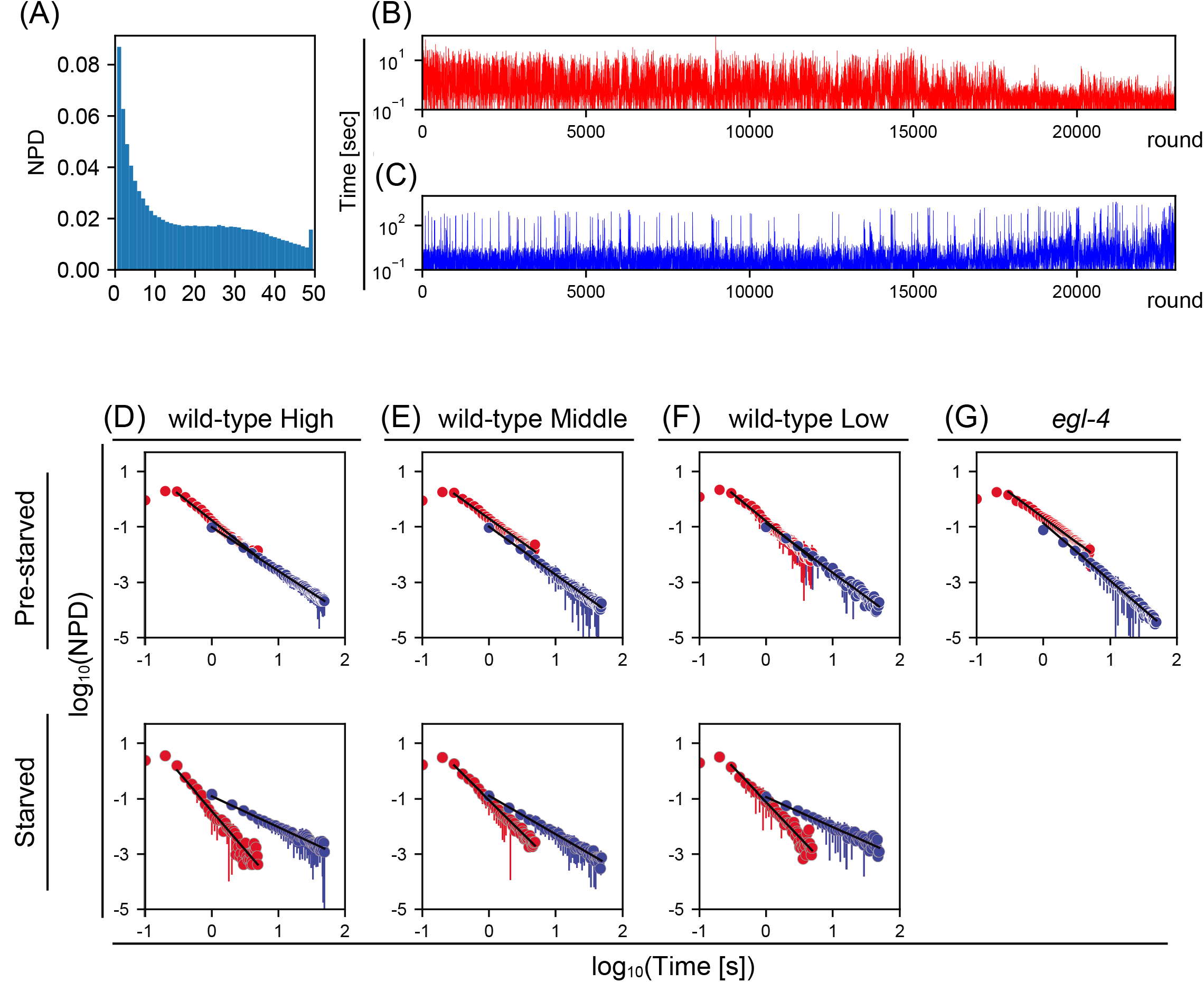
A power law distribution of active/inactive state residence times in *C. elegans* episodic swimming. (A) Appearance frequency of number of active pixels in a representative animal for a single image difference frame. Bimodal distribution of active pixels in normalized probability density (NPD) was separated at 10~20 pixels/frame. The active state threshold was ≥12 active pixels for round series data analysis (details in Methods). (B and C) Round series of residence times in the active (B) and inactive (C) states with the y-axis in log scale. Activity periods above and below the active state threshold (separated by a red horizontal line on 5-s scale in Fig. 2C) were defined as active and inactive periods, respectively. (D-G) Mean and standard deviations of NPD of residence times in active state (red) and in inactive state (blue) among individual animals in indicated time regime were shown in log-log plot. Fitting was performed over 0.3–5.0 s for active states and over 1–50 s for inactive states. The fit line is shown as a black line. Pre-starved vs. Starved time regimes are as described in Figure 2 (wild-type) and the main text (*egl-4* mutants).

The probability distributions of residence times in active and inactive states were linear lines in log-log plots or power law distributions (Fig. 3D-F, and Table 1, fit parameter), indicating that the state residence times lack a specific time scale and thus can be described as exhibiting a scale-free property. Comparing the power law exponents between the two states in each activity class in Pre-starved and Starved time regime, we find that the power law relationship slopes are significantly shallower for the inactive state than for the active state (a “shallow slope” power law relationship indicates a relatively even appearance of longer and shorter residence times, Fig. 3D–F, Table 1 fit parameters and p values in Table 1). These results indicate that the mechanisms that regulate the transition from the active to the inactive state and from the inactive to the active state have distinct scale-free kinetics.

### Multifractal analysis of numerically-generated round series data

Long and short bursts of state residence times in the round series data were clustered rather than random (Fig. 3B and C). We employed multifractal detrended fluctuation analysis (MF-DFA, Methods) to study the temporal structure of the residence-time round series data^37,38^. First, time series of white, pink, and Brown noises were numerically-generated by R software tools for fractional Gaussian noise generation^39,40^, and were subjected to MF-DFA. White noise time series is completely uncorrelated, whereas pink and Brown noise time series have long-range correlation. Noise components of cumulatively summed original time series [*f*(*v, s*) in Eq. 1] was estimated in MF-DFA by removing local trends in each scale, *s*, by linear fitting to various time scales (Methods). As the temporal correlation in the time series became longer, the noise component (i.e. deviation from the trend of cumulative sums of deviations from the average) became smoother (Fig. 4A; upper three series). The calculated average noise component [*F*(*q,s*), Eq. 1] was plotted with temporal resolution for observation in *s*. In *F*(*q,s*) versus *s* plots on a log-log scale, *F*(*q,s*) increased linearly with increasing scale *s* (Fig. 4A; left column of lower three graphs), as is indicative of fractal scaling. Additionally, the slope of *F*(*q, s*) versus *s* plot on a log-log scale increased with an increasing length of temporal correlation of the original time series (Fig. 4A; left column of lower three graphs). The *F*(*q,s*) versus *s* plot gave rise to a multifractal spectrum (Methods and Supplemental Information) showing a relationship between *q*-order (local) Hurst exponent (Hölder spectrum *H*(*q*)) vs. *q*-order singularity dimension (singularity spectrum *D*(*q*))^37,38^.

**Figure 4.**
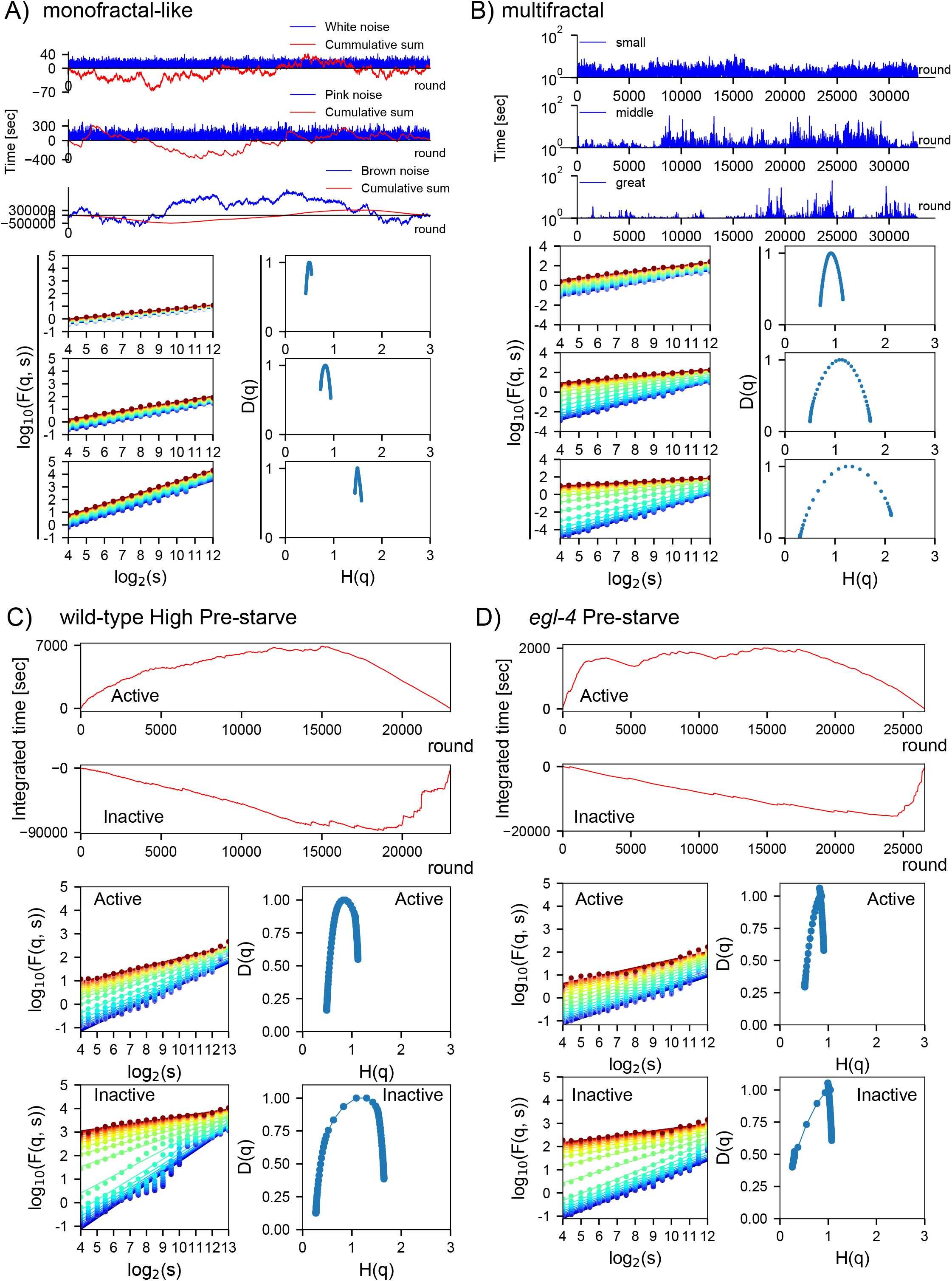
Multifractal analysis of numerically-generated round series and experimentally obtained round series of active and inactive states in a representative *C. elegans* animal. (A-D) MF-DFA of numerically-generated round series with monofractal-like (A) and multifractal (B) properties. Time series with small/middle/great multifractality were generated by the multiplicative cascading processes using log-normal functions whose variances were determined by random noises with small/middle/large variances (^41,42^; see Supplemental Information). Experimentally-obtained Pre-starved active/inactive state round series of High activity wild-type (C) and *egl-4* mutant (D) animals. Time series [blue, in (A) and (B), magnified by arbitrary unit] and cumulative sum of the deviations from the average of values in the time series [red, in (A), (C), and (D)] are shown in upper three (A and B) and two (C and D) graphs. *F*(*q,s*) vs. scale *s* plot data (dots) and their fit functions (lines) are shown at lower *q* values (cold colors) and higher *q* values (hotter colors) in a range of −10 < q < 10 [left column in the lower three (A and B) and two (C and D) graphs]; corresponding multifractal spectra are shown in right columns of each graph. Original white, pink, and Brown noise time series are shown, respectively, after 10, 10^2^, and 10^4^-time magnifications (A).

The singularity dimension is a fractal dimension that represents time series sparsity; a *D*(*q*) < 1 represents sparsely distributed local structures of the entire time series, and a *D*(*q*) approaching 1 representing broadening involvement of the time series. Thereby, the Hurst exponent at *D*(*q*) < 1 represents a Hurst exponent of local structures of time series, which is the local Hurst (Hölder) exponent, whereas the Hurst exponent at *D*(*q*) = 1 represents the Hurst exponent of the entire time series, which we call the global Hurst exponent (*H_peak_*). The Global Hurst exponent *H_peak_* corresponds approximately to the conventional Hurst exponent, which is linked theoretically with the scaling exponent of power spectrum density [*β*, (*S*(*f*)~*f*^-*β*^)], a measure of time series autocorrelation [*β* = 2*H* + 1, (*H* > 0)] (Kiyono, 2015). Therefore, the global Hurst exponent *H_peak_* can be interpreted as an index of behavioral memory in activity time series of behavior. The *H_peak_* values of the cumulative sum of these noise time series were estimated to be approximately 0.5, 1, and 1.5 (Fig. 4A; right column of lower three graphs), consistent with theoretical Hurst exponents of cumulative sums of these noise time series.

Next, multifractal time series were numerically-generated by multiplicative cascading process^41,42^, and were subjected to MF-DFA. The multifractal time series were highly clusterized, such that the variety of amplitudes and durations of temporal clusters became richer with greater multifractality (Fig. 4B; upper three series). Because of the scale-free and self-similar properties of multifractal time series, such temporal clusters are distributed in multiple temporal resolutions. As the variety of amplitudes and durations of temporal clusters increased, the *F*(*q,s*) versus *s* plot slope in a log-log scale varied widely with the exponentiation factor *q* (Fig. 4B; left column of lower three graphs), which enhances or suppresses the noise component from local trends at scale *s* [*f*(*v, s*) in Eq. 1, and Methods]. The *q*-dependent change in slope in a log-log scale *F*(*q, s*) versus *s* plot is indicative of multifractal scaling. In accordance with this *q*-dependent variation, the width of the multifractal spectrum *(width)* widens significantly without a substantial change in *H_peak_* (Fig. 4B; right column of lower three graphs). A wide multifractal spectrum indicates that there are locally clustered structures with various local Hurst exponents sparsely distributed throughout the time series. Therefore, multifractal spectrum *width* in animal activity time series is an index of behavioral complexity. Note that global Hurst exponent *H_peak_* and multifractal spectrum *width* are independently changeable in a scale-free time series with a given power law exponent. See the Supplemental Information for detailed descriptions of numerical generation for noise time series and multifractal analysis, and for an introduction into fractal scaling.

### Episodic swimming is driven by two-state transition with two distinct multifractal kinetics

Global trend in the round series data of active- and inactive-state residence times became gradually shorter and longer, respectively, after transition from the Pre-starved to the Starved time regime (~15,000 rounds; Fig. 3B and C). The noise components of cumulative sums of the deviations from average residence times appeared to differ qualitatively between active and inactive states; that of active states was smoothly curved, whereas that of inactive states was locally straighter in the Pre-starved regime (Fig. 4C).), suggesting that noise properties of active and inactive states may differ. In the log-log scale *F*(*q, s*) versus *s* plot, *F*(*q, s*) in active and inactive states increased almost linearly with *s* and the slope varied with *q* (Fig. 4C; left column of lower two graphs), and the multifractal spectra of active state and inactive states in the Pre-starved time regimes were widely distributed (Fig. 4C; right column of lower two graphs), indicating that the round series of active and inactive states had a multifractal nature.

The Global Hurst exponents *H_peak_* in multifractal spectra of active and inactive states were located around *H*(*q*) =1, indicating that the state residence times have a non-trivial long-range temporal correlation, as is seen in pink noise time series. As shown in Figure 5A-C, before and after starvation in all activity classes, we observed greater *H_peak_* values for the inactive state (average, 1.38) than for the active state (average, 0.85) as well as greater *width* values in the inactive state (average, 1.80) than in the active state (average, 0.64), all p < 0.05). Hence, residence-time round series revealed distinct multifractal properties between active and inactive states, with a longer behavioral memory and greater behavioral complexity for the inactive state than for the active state. We called kinetics that generate scale-free residence times with temporal correlation and temporal clusterization as multifractal kinetics. Accordingly, we were able to explain *C. elegans* episodic swimming with a two-state transition model in which opposite transitions between actively-moving and inactive states are driven by distinct multifractal kinetics (Fig. 6A).

**Figure 5.**
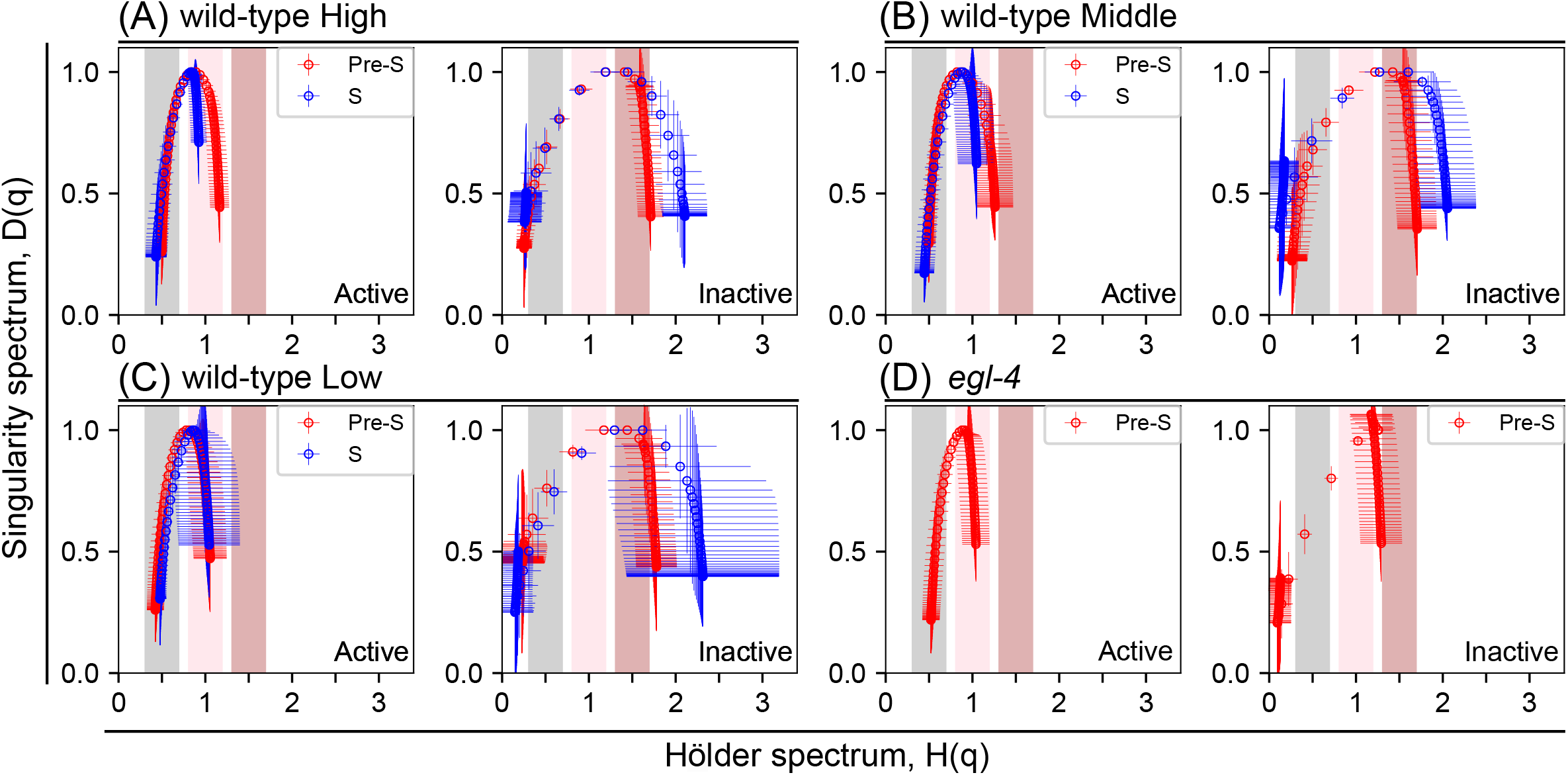
Multifractal spectra averaged from multiple animals. Multifractal spectra means (and standard deviations) were calculated from round series data spectra of active (left) or inactive (right) states from multiple wild-type animals in High (A), Middle (B), and Low (C) activity classes in the Pre-starved (Pre-S, red circles) or Starved (S, blue circles) time regimes. For *egl-4* mutants (D), spectra represent only in Pre-S (red circles) time regime. Note that the spectrum *width* in Pre-starved time regime was as narrow as in Starved time regime in Low activity classes, which was due to too short high activity period during the defined Pre-starved regime among animals in Low activity classes (C).

**Figure 6.**
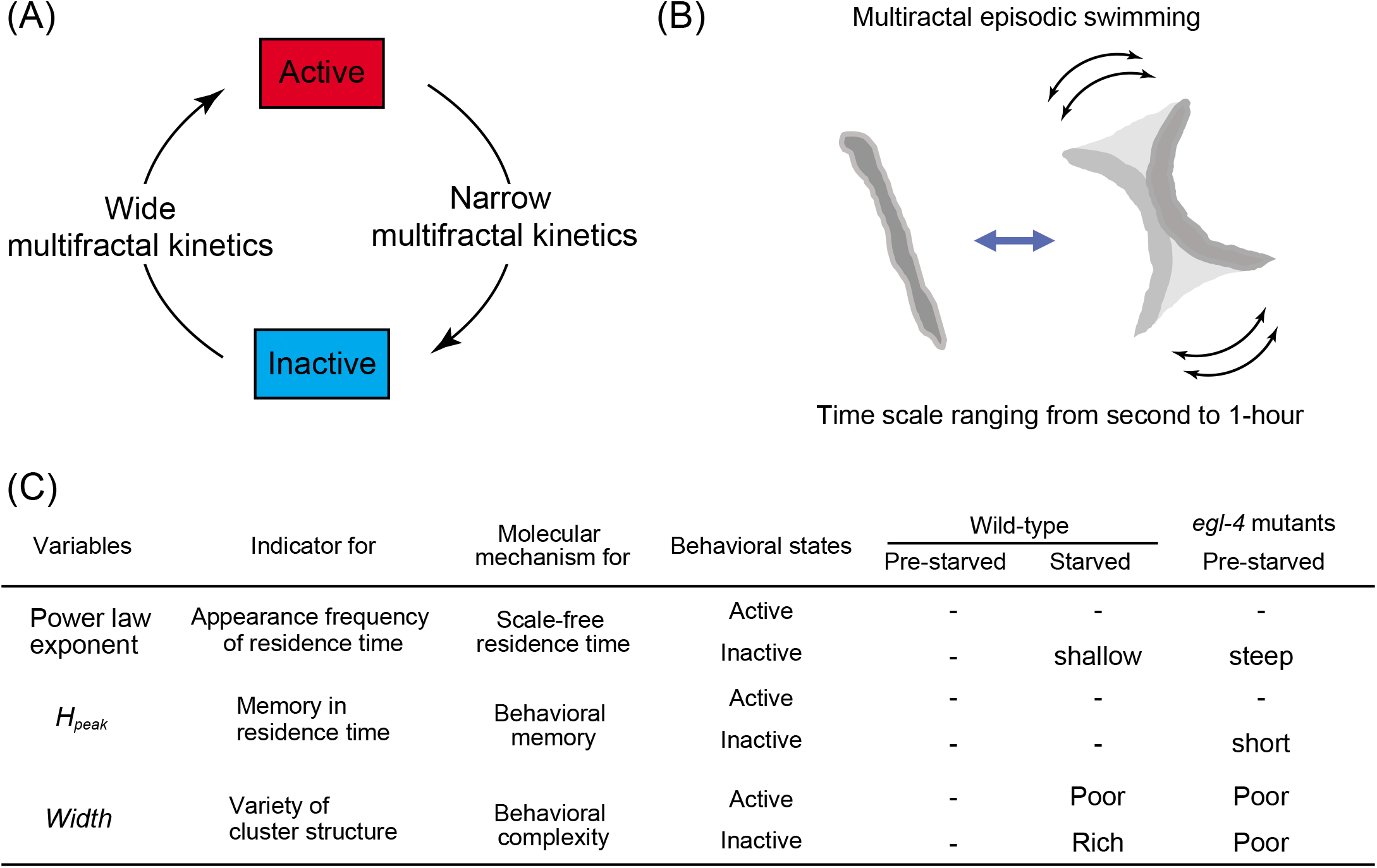
Multifractal episodic swimming in *C. elegans.* (A) Alternate state transitions are driven by distinct kinetics: active-to-inactive and inactive-to-active state transitions follow narrow and wide multifractal kinetics, respectively. (B) Because *C. elegans* episodic swimming is driven by alternating transitions between an actively moving state and an inactive (resting and posing) state over a broad range of temporal scales (i.e. seconds to hours) the residence-time round series data of which are characterized by a multifractal nature, we refer to this behavior as “multifractal episodic swimming”. (C) Summary of our kinetic analyses. See Tables 1 and 2.

The power law exponent showed a more pronounced alteration between in Pre-starved and Starved time regimes in residence-time round series for the inactive state (average, 22.4%) than for the active state (average, 7.0%). The change in the power law exponent was only statistically significant for the inactive state (p values and fit parameters in Table 1). MF-DFA showed that the global Hurst exponent *H_peak_* was not altered significantly by the transition to the Starved time regime in either the active state (average Pre-starved *H_peak_*, 0.86; average Starved *H_peak_*, 0.84) or the inactive state (average Pre-starved *H_peak_*, 1.34; average Starved *H_peak_,* 1.42) *(H_peak_* values, and p values in Table 2, all p > 0.05). However, the multifractal spectrum *width* was altered after starvation, becoming narrower (−17.1%) in the active state and wider (+31.6%) in the inactive state *(width* values, and p values in Table 2, all p < 0.05 except active state in Low class, see Figure 5 legend). These results indicate that food signaling or metabolic state regulates behavior by modulating multifractal kinetics in response to starvation. That is, in response to food starvation, the animals do not simply decrease swimming activity, but show a selective modulation of multifractal kinetics.

### PKG may regulate the multifractal kinetics of *C. elegans* episodic swimming

For molecular dissection of the multifractal kinetics *C. elegans* episodic swimming, we studied the *egl-4(n479)* mutant, in which the temperature sensitive *n479* allele of *egl-4* is associated with a defect in conventional episodic swimming characterized by lengthened continuous swimming periods and a reduced frequency of resting^23,25^. We confirmed that *egl-4(n479)* mutants had reduced frequencies of a persistent inactive state in our quantitative activity time series (Fig. S5B). The *egl-4(n479)* mutants cultured in WormFlo without an energy source were too transparent for body detection in the later portion of the 6-d culture period. This transparency was likely caused by the mutants’ unusually early lipid consumption due to their continuous swimming and lack of long resting; lipids in worm bodies scatter illuminating light enabling imaging as a (relatively) dark object as shown in Figure 2A. Thus high-contrast images of the first tenth of the 6-d culturing period were used for analysis of the Prestarved time regime of *egl-4* mutants.

In *egl-4* mutants, the power law relationships were maintained in active and inactive states, but the power law exponent was preferentially changed in the inactive state (+21.0%) compared to that in active state (+6.9%) (Fig. 3G and D). Power law exponents differed significantly between wild-type worms and *egl-4* mutants only in the inactive state (p-values and fit parameters in Table 1), indicating that the mutant has altered residence times across time scales rather than a defect that is specific to a particular time scale. MF-DFA showed that the shape of the multifractal spectrum in *egl-4* mutants was largely altered in the inactive state (Figs. 4D and 5D). Comparing the global Hurst exponent between *egl-4* mutants and High-activity class wild-type worms, we found that active-state *H_peak_* was similar across the groups (difference, 1.1%), whereas inactive-state *H_peak_* differed significantly between the groups (difference, 18.7%). Meanwhile, spectrum *width* in *egl-4* mutants was reduced significantly relative to that obtained for the High activity wild-type group in both active (−20.9%) and inactive (−14.1%) states *(H_peak_* and *width* values, p values in Table 2). Thus, *egl-4* mutants showed a defect in both temporal memory and temporal clusterization in the inactive state, but showed a defect in only temporal clusterization in the active state. These results are consistent with the possibility that EGL-4/PKG may regulate the multifractal kinetics of behavior, and, more specifically, suggest that EGL-4/PKG may regulate the active- and inactive-state kinetics of *C. elegans* episodic swimming differently.

## Discussion

### *C. elegans* episodic swimming is driven by a multifractal transition cycle

Experimental measurements of temporal changes in biological system variables of interest and the statistical analysis of those data can provide information about the hidden operating principles of that system^43^. *C. elegans* episodic swimming was characterized conventionally by the average residence time of active- and inactive-states^23–25,44^. Here, we applied fractal analysis to study the molecular and genetic mechanisms regulating animal behavior over a 6-d period recorded at a subsecond temporal resolution, showing that episodic swimming is a scale-free process across a 1000-fold range of time scales (Fig. 2C). Because the experimentally-measured average residence times often fail to be determined by values that greatly exceeded the average times, the scale-free process is characterized by a power law exponent in the relationship between appearance frequency and residence time (Supplemental Information). Our multifractal analysis showed long-range memory and local clusterized structures of episodic swimming residence times characterized by a multifractal process. Hence, we referred to the swimming as “multifractal episodic swimming” (Fig. 6B).

A scale-free distribution of actively-moving and inactive residence times has been reported for episodic *Drosophila* behavior^9^; modeling suggested that the scale-free nature of these residence times contributes to maximizing the food exploitation area while optimizing food intake time. Previous analyses have shown long-range temporal correlations in scale-free properties of the behaviors of many species, including fractal (but not multifractal) analysis of *C. elegans* crawling^7,8,10–13^. The fractal nature of *C. elegans* crawling in agar (Alves et al., 2017) is consistent with our long-range memory and clusterization finding. In this study, we extended previous findings by combining a two-state transition model with multifractal analysis. Employing multifractal analysis, we found that scale-free actively-moving and inactive states had a long-range memory and complex local temporal structures. Our analysis indicated that *C. elegans* episodic swimming is characterized by a two-state transition between actively-moving and inactive states, wherein the two transitions are driven by distinct multifractal kinetics. That is, the active-to-inactive transition is driven by a narrow multifractal kinetics (short long-range temporal correlation and low complexity), whereas the inactive-to-active transition is driven by a wide multifractal kinetics (extended long-range temporal correlation and relatively high complexity) (Fig. 6A).

Multifractal episodic motion of *C. elegans* in solid and liquid environments may be an adaptation to food environments. The colonies of bacteria that *C. elegans* worms feed on grow in a fractal shape^45^. In *C. elegans*, the actively-moving state is likely to enable food foraging, whereas the inactive state is likely associated with food intake, egg-laying, or resting to save energy or satiety^24,29–31,46,47^. Temporal correlation in the inactive state gives rise to a series of long and short periods for food intake that may follow the fractally-shaped bacterial colony, whereas temporal correlation in active state round series gives rise to a series of long- and short-distance foraging bouts that may follow the interbranch distances of a fractally-shaped bacterial colony. Thus, the scale-free and temporally structured residence times for food foraging and intake may be adaptive to the fractal shape of bacterial colonies. *C. elegans* survival strategies are altered by food availability. Under starvation conditions, *C. elegans* saves their energy for long-distance foraging and instead spend more effort for balancing resting and food intake at a local area, e.g. through reuptake of their excrements. Food-dependent modulation of multifractal kinetics of behaviors may improve food intake efficiency and reproductive success in natural environments. This possibility should be tested in a modeling study.

Our quantitative studies showed that *egl-4* mutants exhibited different alterations of multifractal kinetics in the active versus the inactive state (Fig. 6C). This result suggests that EGL-4 may regulate the multifractal kinetics of animal behaviors. However, *egl-4* mutants’ defects in behavioral memory and behavioral complexity were smaller in magnitude than the differences between the active and inactive states in wild-type *C. elegans,* and also smaller than the index that characterizes qualitative differences of noise properties among white, pink, and Brown noises. Differences between active and inactive states in wild-type *C. elegans* were comparable to or greater than the index (compare *H_peak_* in Table 2: *H_peak_* of white, pink, and Brown noises differ by 0.5). Therefore, multifractal kinetics in *egl-4* mutants were within the range of kinetics qualitatively same as those in wild-type. In addition, these defects may be caused by a behavioral variation of the *egl-4* mutant strain rather than by loss of function of *egl-4/pkg* per se. It is necessary to apply other fractal analyses and further molecular/genetic studies to examine these possibilities.

### PKG-modulated multifractal transition may be widely applicable to multi-time scale behaviors

Multifractal transition cycle and its PKG-dependent modulation may be shared among many invertebrates. In short-term (sub-minute) observations, *Drosophila*^9^ and *Leptothorax allardycei* worker ants^28^ exhibit bimodal behavioral mode switching^9,28^ similar to that in *C. elegans*. Notably, *Drosophila* residence times in actively-moving and inactive states in a short-term observation period were distributed in a scale-free manner^9^. Interestingly, a bimodal behavioral choice between long-distance moving foragers and dwellers has been shown to be regulated by PKG in *C. elegans*^26,27,48^, *Drosophila*^32^, bees^35^, and ants^36,49^. Both *C. elegans* mutant^26,27^ and *Drosophila* polymorphism^32,50^ local dwellers spend more time on local food intake/resting. Local dwellers conduct brood care in bee hives^35^ and ant nest defense^36,49^. Forager and dweller behavioral phenotypes are switched developmentally in bees, but maintained through the lifespan in the ant caste system. In *C. elegans*^26,27,48^ and ants^36^, long-distance foraging is associated with low PKG activity or expression, whereas local dwelling is associated with high PKG expression. Conversely, long-distance foraging (or local dwelling) is associated with high (low) activity or expression of PKG in flies^32^ and bees^35^. The time scale difference of PKG-active periods between ants and honey bees, and the reversed function of PKG for long-distance foraging or local dwelling between nematodes/ants and flies/bees may reflect a heterochronic evolutionary change in the developmental control of PKG expression^35^, and an evolutionary adaptation of the PKG signaling system in molecular drivers of behavior, respectively. Despite evolutionary changes in time scale and the functional role of PKG, these results indicate that appearance frequency for actively-moving and inactive states is modulated by PKG. Although the role of PKG in the episodic motions of *Drosophila* and ants has not been studied directly, the aforementioned findings suggest that at least episodic motions in *Drosophila* and ants may be regulated by multifractal kinetics and its PKG-dependent modulation.

Surprisingly, PKG-dependent modulation of the transition kinetics in a multifractal transition cycle may be involved in human heartbeat physiology. Electrocardiogram time series^51–53^ of cardiac muscle depolarization and repolarization are characterized by multifractality. The inter-beat (RR) interval is the period between peaks of R waves, which reflect ventricular depolarization. The intra-beat (QT) interval is the period from the peak of Q wave to the end of T wave, which corresponds to the depolarization in the left side of the intraventricular septum and the repolarization of ventricular muscles, respectively. Round series of RR and QT intervals are characterized by a multifractal structure^54^ Interestingly, the shape of the multifractal spectrum for RR intervals differs from that for QT intervals, raising the possibility that QT and RR intervals may be regulated by distinct multifractal kinetics. PKG has been shown to regulate the heartbeat in mice and flies^55–58^. PKG-knockout mice^56^ or cardio-myocyte specific PKG-knockout mice^58^ show hypertension. Although the mechanism is still in dispute^57^, the defects in these knockout mice indicate that PKG is involved in the relaxation phase of the cardiac cycle. Thus, the heartbeat can be regulated by a multifractal transition cycle and is subject to PKG-dependent modulation, similar to that we documented for *C. elegans* episodic swimming. Thus, PKG-modulated multifractal transition cycles may occur in various animal species, including humans, and organs.

### Molecular and system-level mechanisms for multifractal episodic swimming

Based on previously reported models designed to recapitulate scale-free and multifractal time series, we developed hypotheses for molecular and system-level mechanisms underlying the multifractal nature of animal behaviors, and discuss how PKG may be involved in the mechanism. First, the intermittent bursts of a single neuron can be reproduced in chaos dynamical systems^59–65^; owing to their non-Gaussian fluctuation with intermittency, such systems may be associated with multifractality in animal behaviors. Chaotic dynamics reproduced in these models are generated by interactions between fast inflows and slow outflows of ions to/from a neuron. Thus, multifractal nature in behavior might be attributable to chaotic biochemical reactions in a single neuron. Second, when a system is in a certain state, called the criticality, local interactions among system components related to order and noise for randomization of the order reach a critical balance and, eventually, self-similar dynamics emerge spontaneously in the system^66,67^; scale-free dynamics due to the criticality can be associated with multifractality in animal behaviors^68,69^. Experimentally observed scale-free neural activity in intact brains was recapitulated in self-organized criticality-based models that assumed global interaction of neuronal signals in a hierarchically structured neural network representative of brain structures in *C. elegans* and humans^70,71^. Therefore, multifractality in animal behaviors may be attributed to criticality in the hierarchal structure of brains. Third, although implications of molecular and physiological mechanisms in the multifractal nature of animal behaviors have not been reported, the multiplicative cascade model is a simple model that generates multifractal time series (Fig. 4B, Supplemental Information)^41,42,72,73^. Generally, multiplicative cascade models generate multifractal time series via a series of multiplications of log-normal random noise in a cascade manner, enhancing time series variance greatly as it progresses (see Supplemental Information). It has been shown that waiting time distributions of neuronal action potentials exhibit a log-normal distribution consequent to the nonlinearity of reaction systems in neurons^74–76^, whereas hierarchal multiplication may correspond to a cascading signal relay in the brain. Thus, multifractal nature in behaviors may be attributed to both non-linear reactions within neurons and hierarchal structures of neural networks. To reveal molecular- and system-level mechanisms of *C. elegans* multifractal episodic swimming, it will be critical to identify the operating principles that are functioning in multifractal episodic swimming in *C. elegans*. With regard to PKG, selective ectopic expression of PKG in R3 and R4d ring neurons is sufficient to restore behavioral defects in *Drosophila* with *pkg*/*foraging* alleles^77,78^. Additionally, differential PKG expression was identified in a set of five specific neurons between different castes of ants^36^. In *C. elegans,* PKG functions in a limited number of neurons^25^. PKG has been shown to regulate synaptic vesicle cycling, Ca^2+^ influx via G-protein signaling, and axon guidance in certain neuron types^77,79–82^. The endogenous and ectopic expression of PKG in specific neurons and functional analysis of PKG suggest that PKG modulates multifractal kinetics of animal behaviors in single neurons or small numbers of neurons. In this study, our experiments revealed a basic kinetic mechanism of *C. elegans* multifractal episodic swimming. The findings are applicable to diverse fields of interest, including human heartbeat physiology. Due to the wide variety of molecular and genetic tools available for *C. elegans* research, our observation and analysis approach may be used to help reveal conserved mechanisms underlying the multifractal nature of animal physiology and behavior.

## Methods

### *C. elegans* strains used and their maintenance

The Bristol N2 strain was used as wild-type *C. elegans.* N2 and *egl-4(n479)* mutant animals were maintained on agar plates with *E. coli* OP50 strain at 15 °C^83^.

### Design, fabrication, and characterization of microfluidic device

To maximize the number of animals monitored in the recording area, our device employs a two-vertical-compartment structure with the array of culture chambers located over a flow path, whose boundary was partitioned by a porous membrane (Whatman 111115, Nuclepore Hydrophilic Membrane, 10-μm pores, GE Healthcare, USA) (Fig. 1A and B). The upper polydimethylsiloxane (PDMS) chip consisted of a 108-chamber array (1 worm/chamber) with buffer-inlet and exchange solution-outlet ports in the chambers. The lower PDMS chip consisted of a snake-shaped microchannel for supplying liquid buffer to the culture chambers. PDMS chips were fabricated by conventional replica molding with the SU-8 epoxy-based photoresist^84,85^. Animalloading ports (100 μm wide) on the top of each of chamber in the upper PDMS chip were made with a Zing 16 laser (Epilog Laser, Japan). Before culturing and buffer exchanges, the loading ports were sealed with a PDMS sheet. Each culture chamber (2-mm diameter and 0.3-mm height) in the upper PDMS chip was approximately 2 fold-longer and 3 fold-thicker than the ~1-mm-long and ~0.1-mm-wide *C. elegans* body (Fig. 1C). The serpentine buffer supply microchannel in the lower PDMS chip had a 2.2-mm diameter, covering each upper-chip culture chamber. The three parts (upper and lower PDMS chips, and the microporous membrane) were assembled by covalent bonding with an aminosilane coupling agent and oxygen plasma treatment^86^.

### Culturing *C. elegans* in a microfluidic device

Animals at the young adult stage were collected from an agar plate. After an agar plate-culture period (2 d at 24 °C for wild-type and 3 d at 15 °C for *egl-4* mutants), the animals were collected manually in room-temperature M9 buffer and introduced into the microfluidic device by manual pipetting via chamber loading ports (Fig, 1C). With previous devices, individual *C. elegans* were held in a clump structure and released into the chamber by applying deforming high pressure^87–89^. We instead used manual pipetting to avoid applying mechanical stress during loading. To maintain a constant chemical environment in the culture chambers, M9 buffer (with or without 1 g/L glucose and 5 mg/L cholesterol) was perfused continuously with a peristatic pump (Fig. 1B and D) at a flow rate at 5 ml/h. Flow rate was measured by monitoring weight changes in the water discharged from the outlet (data not shown). Buffer exchange in chambers with the flow passing through the porous membrane was confirmed in experiments with a fluorescent solution (Fig. S2). Air bubbles in the buffer supply tube were removed with polytetrafluoroethylene membrane in an Omnifit bubble trap (006BT, Diba Industries Inc., USA) (Fig. 1D).

### Observation of *C. elegans* swimming

*C. elegans* behaviors were recorded at 20 frames/s through a macroscope with an apochromat objective lens (1 ×) (Z16 APO, Leica, Germany) and a CCD camera (1940 × 1460 pixels, 2.8 Megapixel) with a USB3.0 connection (MD028MU-SY, Ximea, Germany). The camera was controlled by Micromanager (https://micro-manager.org). Movies were compressed (H265 codec in FFmpeg) every 10,000 frames (500 s) into mp4 files. *C. elegans* are sensitive to temperature change of 4 °C^90^ and highly sensitive to blue-ultraviolet light, but not to green/yellow light (>545 nm)^91,92^. Temperature of culture chambers on WormFlo were maintained by submerging WormFlo in M9 buffer in a 15-cm-diameter glass dish, whose temperature was maintained by temperature-controlled water supplied from a high-precision water bath (HAAKE, Germany) (Fig. 1D and Fig. S3A and B). A temperature logger (TC-08, Pico Technology, UK) confirmed that the temperature of the M9 buffer in the glass dish was maintained within ±0.5 °C during the 6-day recording period (Fig. S3C). The recording system was covered by a light shield to prevent illumination changes from light fluctuations related to daily lab activities (Fig. 1D). Blue light of illumination light from a halogen lamp was filtered out with a 0.5-mm-thick orange acryl plate (Fig. 1D and S3A); spectrophotometry (USB400, Ocean Optics, USA) confirmed that wavelengths <500 nm were filtered out (Fig. S3D). Standard deviation of light intensity change at the culture chambers during observation period was kept within 10% of the average (Fig. S3E). This culturing and recording system allowed us to monitor individually cultured *C. elegans* in an environment with minimized chemical, light, and temperature perturbations.

### Quantification of *C. elegans* swimming activity

We measured *C. elegans* swimming activity by counting the number of pixels with an intensity over a certain threshold in a matrix obtained from the difference between intensities obtained at t and t + 1 (we referred to it as image difference) using the Open CV module in Python2.7. When an animal moves at a frame interval, the dark pixels detecting its body in image[t] become brighter in image[t +1], yielding an increased image difference (= image[t +1] – image[t]) (Fig. 2A). For pixel counting, the image difference matrix values were in the range of – 256 to +256 at 1940 × 1460 pixels. The pixels with intensity differences greater than 12 value within this range thus exhibit an image difference and are counted as “active” pixels (note that the pixel intensity threshold of 12 is different from the activity threshold (12 pixels/frame) to define residence time in active- and inactive states (Fig. 3A)). The pixel intensity threshold of 12 was determined empirically so that actual movements of animals are detected efficiently while avoiding artefact-based false hits due to thermal noise in a pixel on the camera sensor. Based on pixel counts, the size of the area the animal moved through during each time frame (50 ms) was measured (Fig. 2A). Image and data processing, including compensation for artifactual activity in the time series are described in the Supplemental Information (Figs. S4 and S6). To classify swimming activity, we measured <u>a</u>verage <u>a</u>ctivities during the early half of the recording period (day 0 to day 3; abbreviated “AA0-3”) and the ratio of AA0-3 to <u>a</u>verage <u>a</u>ctivities during the latter half of the recording period (day 3 to day 6; abbreviated “AA3-6”). Animals with high AA0-3 and a low AA0-3 to AA3-6 ratio were classified as High activity. Animals with high AA0-3 and a high AA0-3 to AA3-6 ratio were classified as Middle activity. Animals with low AA0-3 and a low AA0-3 to AA3-6 ratio were classified as Low activity. The threshold of high AA0-3 was set empirically at 0.75 and the threshold of a high ratio was set to 0.01. Note that 0.75 threshold values are low due to the bursty/sparse nature of *C. elegans* swimming activity.

### Data analysis

Active-state residence time was defined as the period after the start of the activity burst to the end of the burst; inactive-state residence time was defined as the period starting immediately after the end of activity burst to the next round of activity burst in the activity time series. Alternating-state round series data were obtained by thresholding time series of swimming activity at 12 pixels/frame, which corresponded to the valley of a bimodal activity distribution (Fig. 3A).

MF-DFA was performed in Python software^93^ using

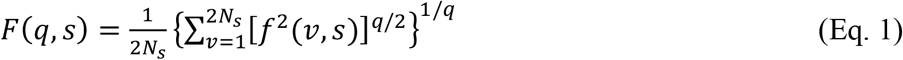

where *f*(*v, s*) is the noise component or fluctuations from the local trend of cumulative sums of the deviation of residence time from the average residence times in the *v*^th^ segment at the temporal resolution for observation (s). MF-DFA was derived from detrended fluctuation analysis (DFA)^94^; the equation used in DFA corresponds to the equation used in MF-DFA when *q* = 2 in Eq. 1). In operation 1, the entire cumulative sum series was segmented into *Ns* segments at scale *s*, and the local trend in each *v*^th^ segment at scale *s* was determined by piecewise fitting with a linear function. In operation 2, the amplitude of fluctuations from the local linear trend at each segment was enhanced (or suppressed) to a large (or small) amplitude of *f*(*v,s*) by exponentiating with positive (or negative) *q*-values; *q* value-exponentiated *f*(*v,s*) was summed over all the *v*^th^ segments. In operation 3. the operations 1 and 2 above were performed bidirectionally, forward and backward, on the round series (in total 2 × *N_s_* segments) to obtain *F*(*q, s*). *F*(*q, s*) versus *s* plots were log-log plotted, and each *q*-value was fit with a linear function to obtain a local Hurst (Hólder) exponent. Finally, multifractal spectrum [*q*-order (local) Hurst exponent, or Hólder spectrum *H*(*q*) vs. *q*-order singularity dimension, or singularity spectrum *D*(*q*)] was obtained from *F*(*q,s*) versus *s* at each *q*-value by Legendre transformation. Linear fitting to data in a log-log plot minimizes relative error between the fit function (y_fit) and the data (y_data), which is log(y_fit/y_data) (= log(y_fit)- log(y_data)). Linear fitting to data in a log-log plot avoids biased fitting due to errors being weighted by high-value data points that happens when fitting is performed by minimizing absolute error (y_fit – y_data). Animals whose chambers had long-term retained bubbles, and wild-type animals which were transparent at the final movie frame were eliminated from the data analysis.

### Statistical analysis

Student t-tests and chi-squared tests for independence were performed with scipy.stats.chi2_contingency and scipy.stats.ttest_ind, respectively^95^.

## Author contributions

Conceptualization: Yukinobu Arata and Hiroaki Takagi

Formal analysis: Peter Jurica and Yukinobu Arata,

Funding acquisition: Yukinobu Arata and Yasushi Sako

Investigation: Yusaku Ikeda, Yukinobu Arata, and Peter Jurica

Methodology: Yusaku Ikeda and Hiroshi Kimura

Writing: original draft: Yukinobu Arata

Writing – review & editing: Hiroaki Takagi, Struzik Zbigniew, Ken Kiyono, and Yasushi Sako

The authors declare no conflict of interest.

This article contains supporting information online at

This open access article is distributed under Creative Commons Attribution License (CC BY).

## Data deposition

The *C. elegans* swimming activity time series and movie data reported in this paper have been deposited in Systems Science of Biological Dynamics (SSBD) database ^1^, http://ssbd.qbic.riken.jp/set/20190401/

## Abbreviations

(PKG): cGMP-dependent kinase

## Acknowledgements

We thank the *Caenorhabditis* Genetics Center (CGC) for *C. elegans* strains; the CGC is funded by NIH Office of Research Infrastructure Programs (P40 OD010440). We are grateful to Yuki Shindo for providing valuable help with Python programing, to Seiichi Uchida at Kyushu University for sharing computational power for recording, and Hideitsu Hino at The Institute of Statistical Mathematics for advices for statistical tests. We are grateful to RStudio Team (2018) for RStudio: Integrated Development for R. RStudio, Inc., Boston, MA URL http://www.rstudio.com/.

## References

1 Tohsato, Y., Ho, K. H., Kyoda, K. & Onami, S. SSBD: a database of quantitative data of spatiotemporal dynamics of biological phenomena. Bioinformatics 32, 3471–3479, doi:10.1093/bioinformatics/btw417 (2016).

2 Reddy, A. B. & Rey, G. Metabolic and nontranscriptional circadian clocks: eukaryotes. Annu Rev Biochem 83, 165–189, doi:10.1146/annurev-biochem-060713-035623 (2014).

3 Mellor, J. The molecular basis of metabolic cycles and their relationship to circadian rhythms. Nat Struct Mol Biol 23, 1035–1044 (2016).

4 Iannaccone, P. M. & Khokha, M. Fractal Geometry in Biological Systems: An Analytical Approach. (CRC Press, 1996).

5 Mandelbrot, B. B. The fractal geometry of nature. Updated and augm. edn, (W.H. Freeman, 1983).

6 Bunde, A. & Havlin, S. Fractals in Science. (Springer-Verlag, 1994).

7 Kembro, J. M., Flesia, A. G., Gleiser, R. M., Perillo, M. A. & Marin, R. H. Assessment of long-range correlation in animal behavior time series: The temporal pattern of locomotor activity of Japanese quail *(Coturnix coturnix)* and mosquito larva *(Culex quinquefasciatus)*. Physica A 392, 6400–6413 (2013).

8 Guzman, D. A. et al. The fractal organization of ultradian rhythms in avian behavior. Sci Rep 7, 684, doi:10.1038/s41598-017-00743-2 (2017).

9 Cole, B. J. Fractal Time in Animal Behavior - the Movement Activity of *Drosophila*. Anim Behav 50, 1317–1324 (1995).

10 Alves, L. G. A. et al. Long-range correlations and fractal dynamics in *C. elegans*: Changes with aging and stress. Phys Rev E 96 (2017).

11 Haris, K., Chakraborty, B., Menezes, A., Sreepada, R. A. & Fernandes, W. A. Multifractal detrended fluctuation analysis to characterize phase couplings in seahorse (*Hippocampus kuda*) feeding clicks. J Acoust Soc Am 136, 1972–1981, doi:10.1121/1.4895713 (2014).

12 Seuront L, M.C., B. & J.R., S. in Handbook of Scaling Methods in Aquatic Ecology: Measurement, Analysis, Simulation (eds Laurent Seuront & Peter G. Strutton) 333 (CRC Press, 2003).

13 Seuront, L., Schmitt, F. G., Brewer, M. C., Strickler, J. R. & Souissi, S. From random walk to multifractal random walk in zooplankton swimming behavior. Zool Stud 43, 498–510 (2004).

14 Hu, K., Van Someren, E. J., Shea, S. A. & Scheer, F. A. Reduction of scale invariance of activity fluctuations with aging and Alzheimer’s disease: Involvement of the circadian pacemaker. Proc Natl Acad Sci USA 106, 2490–2494, doi:10.1073/pnas.0806087106 (2009).

15 Hausdorff, J. M. et al. Altered fractal dynamics of gait: reduced stride-interval correlations with aging and Huntington’s disease. J Appl Physiol (1985) 82, 262–269, doi:10.1152/jappl.1997.82.1.262 (1997).

16 Hausdorff, J. M. Gait dynamics, fractals and falls: finding meaning in the stride-to-stride fluctuations of human walking. Hum Mov Sci 26, 555–589, doi:10.1016/j.humov.2007.05.003 (2007).

17 Kobayashi, M. & Musha, T. 1/f fluctuation of heartbeat period. IEEE Trans Biomed Eng 29, 456–457, doi:10.1109/TBME.1982.324972 (1982).

18 Ivanov, P. C. et al. Scaling behaviour of heartbeat intervals obtained by wavelet-based time-series analysis. Nature 383, 323–327, doi:10.1038/383323a0 (1996).

19 Stam, C. J. & de Bruin, E. A. Scale-free dynamics of global functional connectivity in the human brain. Hum Brain Mapp 22, 97–109, doi:10.1002/hbm.20016 (2004).

20 Linkenkaer-Hansen, K., Nikouline, V. V., Palva, J. M. & Ilmoniemi, R. J. Long-range temporal correlations and scaling behavior in human brain oscillations. J Neurosci 21, 1370–1377 (2001).

21 Struzik, Z. R., Hayano, J., Sakata, S., Kwak, S. & Yamamoto, Y. *1/f* scaling in heart rate requires antagonistic autonomic control. Phys Rev E Stat Nonlin Soft Matter Phys 70, 050901, doi:10.1103/PhysRevE.70.050901 (2004).

22 Goldberger, A. L. et al. Fractal dynamics in physiology: alterations with disease and aging. Proc Natl AcadSci USA 99 Suppl 1, 2466–2472, doi:10.1073/pnas.012579499 (2002).

23 Ghosh, R. & Emmons, S. W. Episodic swimming behavior in the nematode *C. elegans*. J Exp Biol 211, 3703–3711, doi:10.1242/jeb.023606 (2008).

24 McCloskey, R. J., Fouad, A. D., Churgin, M. A. & Fang-Yen, C. Food responsiveness regulates episodic behavioral states in *Caenorhabditis elegans*. J Neurophysiol 117, 1911–1934, doi:10.1152/jn.00555.2016 (2017).

25 Ghosh, R. & Emmons, S. W. Calcineurin and protein kinase G regulate *C. elegans* behavioral quiescence during locomotion in liquid. BMC Genet 11, 7, doi:10.1186/1471-2156-11-7 (2010).

26 Fujiwara, M., Sengupta, P. & McIntire, S. L. Regulation of body size and behavioral state of *C. elegans* by sensory perception and the EGL-4 cGMP-dependent protein kinase. Neuron 36, 1091–1102 (2002).

27 L’Etoile, N. D. et al. The cyclic GMP-dependent protein kinase EGL-4 regulates olfactory adaptation in C. elegans. Neuron 36, 1079–1089 (2002).

28 Cole, B. J. Short-Term Activity Cycles in Ants - Generation of Periodicity by Worker Interaction. Am Nat 137, 244–259, doi:10.1086/285156 (1991).

29 McConnell, M. W. & Fitzpatrick, M. J. ‘Foraging’ for a place to lay eggs: A genetic link between foraging behaviour and oviposition preferences. Plos One 12 (2017).

30 Sokolowski, M. B. Social interactions in “simple” model systems. Neuron 65, 780–794, doi:10.1016/j.neuron.2010.03.007 (2010).

31 Reaume, C. J. & Sokolowski, M. B. cGMP-dependent protein kinase as a modifier of behaviour. Handb Exp Pharmacol, 423–443, doi: 10.1007/978-3-540-68964-5_18 (2009).

32 Osborne, K. A. et al. Natural behavior polymorphism due to a cGMP-dependent protein kinase of *Drosophila*. Science 277, 834–836, doi:DOI 10.1126/science.277.5327.834 (1997).

33 Sokolowski, M. B. Foraging Strategies of Drosophila-Melanogaster - a Chromosomal Analysis. Behav Genet 10, 291–302 (1980).

34 Kaun, K. R. et al. Natural variation in food acquisition mediated via a *Drosophila* cGMP-dependent protein kinase. J Exp Biol 210, 3547–3558, doi:10.1242/jeb.006924 (2007).

35 Ben-Shahar, Y., Robichon, A., Sokolowski, M. B. & Robinson, G. E. Influence of gene action across different time scales on behavior. Science 296, 741–744, doi:10.1126/science.1069911 (2002).

36 Lucas, C. & Sokolowski, M. B. Molecular basis for changes in behavioral state in ant social behaviors. Proc Natl Acad Sci U S A 106, 6351–6356, doi:10.1073/pnas.0809463106 (2009).

37 Kantelhardt, J. W. et al. Multifractal detrended fluctuation analysis of nonstationary time series. Physica A 316, 87–114 (2002).

38 Ihlen, E. A. Introduction to multifractal detrended fluctuation analysis in matlab. Front Physiol 3, 141, doi:10.3389/fphys.2012.00141 (2012).

39 Beran, J., Whitcher, B. & Maechler, M. longmemo; Statistics for Long-Memory Processes (Book Jan Beran), and Related Functionality, R package Version 1.1-1. (2018).

40 Ihaka, R. & Gentleman, R. R: A Language for Data Analysis and Graphics. Journal of Computational and Graphical Statistics 5, 299–314, doi:10.2307/1390807 (1996).

41 Bacry, E., Delour, J. & Muzy, J. R. Multifractal random walk. Phys Rev E 64 (2001).

42 Kiyono, K., Struzik, Z. R. & Yamamoto, Y. Estimator of a non-Gaussian parameter in multiplicative lognormal models. Phys Rev E 76 (2007).

43 Arata, Y. & Takagi, H. Quantitative Studies for Cell-Division Cycle Control. Frontiers in Physiology 10, doi:10.3389/fphys.2019.01022 (2019).

44 Gonzales, D. L., Zhou, J., Fan, B. & Robinson, J. T. A microfluidic-induced C. elegans sleep state. Nat Commun 10, 5035, doi:10.1038/s41467-019-13008-5 (2019).

45 Matsuyama, T. & Matsushita, M. Fractal morphogenesis by a bacterial cell population. Crit Rev Microbiol 19, 117–135, doi:10.3109/10408419309113526 (1993).

46 You, Y. J., Kim, J., Raizen, D. M. & Avery, L. Insulin, cGMP, and TGF-beta signals regulate food intake and quiescence in C-elegans: A model for satiety. Cell Metab 7, 249–257 (2008).

47 Raizen, D. M. et al. Lethargus is a Caenorhabditis elegans sleep-like state. Nature 451, 569–U566 (2008).

48 Raizen, D. M., Cullison, K. M., Pack, A. I. & Sundaram, M. V. A novel gain-of-function mutant of the cyclic GMP-dependent protein kinase egl-4 affects multiple physiological processes in Caenorhabditis elegans. Genetics 173, 177–187, doi:10.1534/genetics.106.057380 (2006).

49 Ingram, K. K., Oefner, P. & Gordon, D. M. Task-specific expression of the foraging gene in harvester ants. Mol Ecol 14, 813–818, doi:10.1111/j.1365-294X.2005.02450.x (2005).

50 Osborne, K. A., de Belle, J. S. & Sokolowski, M. B. Foraging behaviour in Drosophila larvae: mushroom body ablation. Chem Senses 26, 223–230, doi:10.1093/chemse/26.2.223 (2001).

51 Morhrman, D. E. & Heller, L. J. Cardiovascular Physiology. 7th edn, (McGraw-Hill Medical, 2010).

52 Lilly, L. S. Pathophysiology of Heart Disease: A Collaborative Project of Medical Students and Faculty. (Wolters Kluwer Health, 2016).

53 Dubin, D. Rapid Interpretation of EKGs. 6th edn, (Cover Pubulishing Co, 1998).

54 Lewis, M. J., Short, A. L. & Suckling, J. Multifractal characterisation of electrocardiographic RR and QT time-series before and after progressive exercise. Comput Methods Programs Biomed 108, 176–185, doi:10.1016/j.cmpb.2012.02.014 (2012).

55 Johnson, E., Sherry, T., Ringo, J. & Dowse, H. Modulation of the cardiac pacemaker of *Drosophila:* cellular mechanisms. J Comp Physiol B 172, 227–236, doi:10.1007/s00360-001-0246-8 (2002).

56 Pfeifer, A. et al. Defective smooth muscle regulation in cGMP kinase I-deficient mice. EMBO J 17, 3045–3051, doi:10.1093/emboj/17.11.3045 (1998).

57 Feil, R., Lohmann, S. M., de Jonge, H., Walter, U. & Hofmann, F. Cyclic GMP-dependent protein kinases and the cardiovascular system: insights from genetically modified mice. Circ Res 93, 907–916, doi:10.1161/01.RES.0000100390.68771.CC (2003).

58 Wegener, J. W. et al. cGMP-dependent protein kinase I mediates the negative inotropic effect of cGMP in the murine myocardium. Circ Res 90, 18–20 (2002).

59 Fan, Y. S. & Holden, A. V. Bifurcations, Burstings, Chaos and Crises in the Rose-Hindmarsh Model for Neuronal-Activity. Chaos Soliton Fract 3, 439–449 (1993).

60 Gu, H. G. & Xiao, W. W. Difference Between Intermittent Chaotic Bursting and Spiking of Neural Firing Patterns. Int J Bifurcat Chaos 24 (2014).

61 Izhikevich, E. M. Neural excitability, spiking and bursting. Int J Bifurcat Chaos 10, 1171–1266 (2000).

62 Chay, T. R. Chaos in a 3-Variable Model of an Excitable Cell. Physica D 16, 233–242 (1985).

63 Fan, Y. S. & Chay, T. R. Generation of periodic and chaotic bursting in an excitable cell model. Biol Cybern 71, 417–431 (1994).

64 Holden, A. V. & Fan, Y. From Simple to Complex Oscillatory Behaviour via Intermittent Chaos in the Rose-Hindmarsh Model For Neuronal Activity. Chaos, Solitons & Fractals 2, 349–369, doi:https://doi.org/10.1016/0960-0779(92)90012-C (1992).

65 Canavier, C. C., Clark, J. W. & Byrne, J. H. Routes to chaos in a model of a bursting neuron. Biophys J 57, 1245–1251, doi:10.1016/S0006-3495(90)82643-6 (1990).

66 Beggs, J. M. The criticality hypothesis: how local cortical networks might optimize information processing. Philos T R Soc A 366, 329–343 (2008).

67 Bak, P., Tang, C. & Wiesenfeld, K. Self-organized criticality: An explanation of the 1/f noise. Phys Rev Lett 59, 381–384, doi:10.1103/PhysRevLett.59.381 (1987).

68 Schertzer, D. & Lovejoy, S. Multifractal Generation of Self-Organized Criticality. Ifip Trans A 41, 325–339 (1994).

69 Tebaldi, C., De Menech, M. & Stella, A. L. Multifractal scaling in the Bak-Tang-Wiesenfeld sandpile and edge events. Physical Review Letters 83, 3952–3955 (1999).

70 Rubinov, M., Sporns, O., Thivierge, J. P. & Breakspear, M. Neurobiologically realistic determinants of self-organized criticality in networks of spiking neurons. PLoS Comput Biol 7, e1002038, doi:10.1371/journal.pcbi.1002038 (2011).

71 Moretti, P. & Munoz, M. A. Griffiths phases and the stretching of criticality in brain networks. Nat Commun 4, 2521, doi:10.1038/ncomms3521 (2013).

72 Arneodo, A., Manneville, S., Muzy, J. F. & Roux, S. G. Revealing a lognormal cascading process in turbulent velocity statistics with wavelet analysis. Philos TR Soc A 357, 2415–2438 (1999).

73 Meneveau, C. & Sreenivasan, K. R. The Multifractal Nature of Turbulent Energy-Dissipation. J Fluid Mech 224, 429–484 (1991).

74 Petersen, P. C. & Berg, R. W. Lognormal firing rate distribution reveals prominent fluctuation-driven regime in spinal motor networks. Elife 5, doi:10.7554/eLife.18805 (2016).

75 Buzsaki, G. & Mizuseki, K. The log-dynamic brain: how skewed distributions affect network operations. Nat Rev Neurosci 15, 264–278 (2014).

76 Roxin, A., Brunel, N., Hansel, D., Mongillo, G. & van Vreeswijk, C. On the distribution of firing rates in networks of cortical neurons. J Neurosci 31, 16217–16226, doi:10.1523/JNEUROSCI.1677-11.2011 (2011).

77 Peng, Q. et al. cGMP-Dependent Protein Kinase Encoded by *foraging* Regulates Motor Axon Guidance in *Drosophila* by Suppressing Lola Function. J Neurosci 36, 4635–4646, doi:10.1523/JNEUROSCI.3726-15.2016 (2016).

78 Kuntz, S., Poeck, B., Sokolowski, M. B. & Strauss, R. The visual orientation memory of *Drosophila* requires Foraging (PKG) upstream of Ignorant (RSK2) in ring neurons of the central complex. Learn Mem 19, 337–340, doi:10.1101/lm.026369.112 (2012).

79 Eguchi, K., Nakanishi, S., Takagi, H., Taoufiq, Z. & Takahashi, T. Maturation of a PKG-dependent retrograde mechanism for exoendocytic coupling of synaptic vesicles. Neuron 74, 517–529, doi:10.1016/j.neuron.2012.03.028 (2012).

80 Wang, X. & Robinson, P. J. Cyclic GMP-dependent protein kinase and cellular signaling in the nervous system. J Neurochem 68, 443–456, doi:10.1046/j.1471-4159.1997.68020443.x (1997).

81 Dason, J. S., Allen, A. M., Vasquez, O. E. & Sokolowski, M. B. Distinct functions of a cGMP-dependent protein kinase in nerve terminal growth and synaptic vesicle cycling. J Cell Sci 132, doi:10.1242/jcs.227165 (2019).

82 Krzyzanowski, M. C. et al. The C. elegans cGMP-dependent protein kinase EGL-4 regulates nociceptive behavioral sensitivity. PLoS Genet 9, e1003619, doi:10.1371/journal.pgen.1003619 (2013).

83 Brenner, S. The genetics of Caenorhabditis elegans. Genetics 77, 71–94 (1974).

84 Lorenz, H. et al. High-aspect-ratio, ultrathick, negative-tone near-UV photoresist and its applications for MEMS. Sensor Actuat a-Phys 64, 33–39 (1998).

85 Hosokawa, K., Fujii, T. & Endo, I. Handling of picoliter liquid samples in a poly(dimethylsiloxane)-based microfluidic device. Anal Chem 71, 4781–4785 (1999).

86 Kimura, H., Yamamoto, T., Sakai, H., Sakai, Y. & Fujii, T. An integrated microfluidic system for longterm perfusion culture and on-line monitoring of intestinal tissue models. Lab on a Chip 8, 741–746 (2008).

87 Chung, K. et al. Microfluidic chamber arrays for whole-organism behavior-based chemical screening. Lab Chip 11, 3689–3697, doi:10.1039/c1lc20400a (2011).

88 Li, S., Stone, H. A. & Murphy, C. T. A microfluidic device and automatic counting system for the study of *C. elegans* reproductive aging. Lab Chip 15, 524–531, doi:10.1039/c4lc01028k (2015).

89 Hulme, S. E. et al. Lifespan-on-a-chip: microfluidic chambers for performing lifelong observation of C. elegans. Lab Chip 10, 589–597, doi:10.1039/b919265d (2010).

90 Simonetta, S. H., Migliori, M. L., Romanowski, A. & Golombek, D. A. Timing of locomotor activity circadian rhythms in *Caenorhabditis elegans*. PLoS One 4, e7571, doi:10.1371/journal.pone.0007571 (2009).

91 Edwards, S. L. et al. A novel molecular solution for ultraviolet light detection in *Caenorhabditis elegans*. PLoSBiol 6, e198, doi:10.1371/journal.pbio.0060198 (2008).

92 Ward, A., Liu, J., Feng, Z. & Xu, X. Z. Light-sensitive neurons and channels mediate phototaxis in *C. elegans*. Nat Neurosci 11, 916–922, doi:10.1038/nn.2155 (2008).

93 Jurica, P. Multifractal analysis for all. Frontiers in Physiology 6, doi:UNSP 27 10.3389/fphys.2015.00027 (2015).

94 Peng, C. K., Havlin, S., Stanley, H. E. & Goldberger, A. L. Quantification of scaling exponents and crossover phenomena in nonstationary heartbeat time series. Chaos 5, 82–87, doi:10.1063/1.166141 (1995).

95 Jones, E., Oliphant, T., Peterson, P. & others. SciPy: Open Source Scientific Tools for Python [Online; accessed 2019-04-09], <“http://www.scipy.org/“> (2001).

