## Supplemental information for "*C. elegans* episodic swimming is driven by multifractal kinetics"

**Self-similarity and scale-free property**

Fractal patterns are observed in spatial and temporal patterns throughout nature, such as in coastal landscapes, rivers, mountains, clouds, moon craters, and galaxy distributions, as well as in the structures of bacterial colonies, wild plant growth, neuronal systems, blood vessels, and pulmonary airways [1-3]. They are characterized by self-similarity and scale-free, properties that can be demonstrated geometrically in two-dimensional fractal patterns called Koch curves. In Koch curves, an elemental shape is drawn repeatedly over a broad range of spatial-scale magnifications to form a nested self-similar structure (Fig. S1A). Koch curves can continue to expand with increasing resolutions, indicating that their characteristics are independent of length or scale, and that they are thus scale-free property (Fig. S1A). Their structure follows a power law exponent between curve lengths and the spatial resolution of observation. This power law-based scaling relationship represents a fractal dimension. As a Koch curve more fully fills a two-dimensional space, its fractal dimension approaches 2, the dimensionality of a smooth (two-dimensional) surface (Fig. S1A). Thus, self-similar, scale-free spatial patterns are characterized by fractal dimension as a generalization of Euclidian dimensional space.

**The Hurst exponent and local Hurst (Hölder) exponent**

In one dimension (e.g. a temporal time series), self-similarity and the scale-free property are also characterized by a fractal dimension ($D$). The fractal dimension of a one-dimensional time series is theoretically linked with the scaling exponent of power spectrum density ($\beta,\ (S(f){\sim}f^{-\beta}$), an index of autocorrelation of the time series. The fractal dimension of a one-dimensional time series and the scaling exponent of power spectrum density are theoretically equivalent to the Hurst exponent ($H$) under certain conditions ($D = 2 - H$, $(0 < H < 1)$ [2,4] and $\beta = 2H + 1$, $(H > 0)$ [5]). Thus, the Hurst exponent represents the geometrical property of one-dimensional time series as a "memory" of temporal events in a time series. One-dimensional trajectories of Brownian particles, which are self-similar and scale-free (Fig. S1B), highly fill "two-dimensional space" between position-time coordinates, such that the fractal dimension of these trajectories reaches 1.5, exceeding the 1.0 dimension of smooth lines. Long-range temporal correlation in the trajectories of Brownian particles (and the cumulative sum of the trajectories) is characterized by Hurst exponent 0.5 (and 1.5, respectively; in general, the Hurst exponent is increased by one from that of the original time series after taking a cumulative sum) [6,7]. Scale-free time series with non-trivial long-range temporal correlation (e.g. pink noise) in which a positive increment tends to follow a negative increment, and vice versa, fills two-dimensional space extensively such that the fractal dimension of the time series exceeds the dimensionality of smooth lines. The non-trivial long-range correlation in pink noise time series (and the cumulative sum of the time series) are characterized with Hurst exponent 0 (and 1, respectively) [6,7]. Time series without temporal correlation (e.g. white noise) does not have a self-similar structure, preventing the definition of a fractal dimension, whereas cumulative sum of white noise time series is characterized by a Hurst exponent of 0.5 [6,7].

The Hurst exponent was generalized to characterize a one-dimensional time-series of the velocity field of fully-developed turbulence in fluid dynamics due to the multifractality [8]. Turbulences, such as in fast-flowing rivers and billowing storm clouds, encompass interacting various-sized unstable vortices, whose interactions cause drag forces that oppose the surrounding fluid flow. Within an observable temporal resolution, time series of turbulence velocity fields exhibit temporal clusters of various amplitudes and durations, mainly due to stochastic separation of large and small eddies [8]. Temporal clusterization is observed in each of temporal resolutions because the time series has self-similarity. Eventually, the velocity time series will be locally interwoven with a huge variety of temporal clusters. Temporal clusterization of local temporal events with highly variant non-Gaussian fluctuation is a source of time series complexity and is characterized by different local Hurst exponents (a.k.a. Hölder exponents). A multifractal spectrum of a time series shows a relationship between $q$-order (local) Hurst exponent, or Hölder spectrum $H(q)$ vs. $q$-order singularity dimension, or singularity spectrum $D(q)$ [6,7]. The singularity dimension is a fractal dimension that represents sparsity for local data points that have a certain local Hurst exponent in the time series; when singularity dimension is 1 (like a smooth line), the corresponding local Hurst exponent represents the global structure of the time series. When the singularity dimension is less than 1 (between that of a point and a line), the corresponding local Hurst exponents represent sparsely distributed local structures of the time series. A wide multifractal spectrum indicates that locally clusterized structures are sparsely distributed in the time series. Therefore, the width of a multifractal spectrum is a measure of the complexity of temporal clusterization in the time series. Note that long-range temporal correlation and temporal clusterization complexity are independently changeable indexes at a given power law exponent of frequency.

**Numerical generation of noise time series**

To generate the noise time series in Figure 4A, we used the R package "longmemo" for fractional Gaussian noise generation [9,10]. For the white noise time series, we generated uncorrelated and normally distributed random time series with a zero mean. For the Brown noise time series, we generated an integrated series of white noise time series. For the pink noise time series, we generated a time series from a fractional Gaussian process $B(t)$ with $H$ = 0.99 and a zero mean ($Cov(B(t), B(t+\tau)) = 1/2(t^{2H} + (t+\tau)^{2H} - \tau^{2H})$, $0 < \mathrm{H} < 1$, where $H$ is the Hurst exponent) [11]. Absolute white noise and pink noise values are shown in Figure 4A. To generate the multifractal time series in Figure 4B, we employed a multiplicative cascade model, $x_i = \xi_i \exp\left[\sum_{j=1}^{m} \omega^{(j)}\left(\left\lfloor\frac{i-1}{2^{m-j}}\right\rfloor\right)\right]$, where $\{\xi_i\}$ is a pink noise time series with $H$ = 0.99 and zero mean, $\omega^{(j,k)}$ are independent Gaussian random variables, $i$ is time index, and $m$ is the total number of cascade steps [5]. At the first cascade step ($j$ = 1), the pink noise time series $\{\xi_i\}$ $(i = 1,2,3,,,2m)$ was divided into early and late halves of the sub time-regions: the early half ($k$ = 0) $(i = 1,2,3,,, m)$ and the late half ($k$ = 1) $(i = m+1, m+2, ,,, 2m)$. Each half of the time series was multiplied with each of two independent log-normal noise series with length $m$ generated with $\exp\left[\omega^{(1,k)}\right]$. At the second cascade step $(j = 2)$, the early and late time regions in the first cascade step were further divided into early and late sub time-regions ($k$ = 0, 1, 2, and 3, the length of each of the four sub time-regions has a length of $m$/4.). Each of the four sub time-regions was multiplied with each of four independent log-normal noise series with a length of $m$/4 generated by $\exp\left[\omega^{(2,k)}\right]$. At the $j$ cascade step, each time-region in the *j-1* cascade step was again divided into early and late sub time-regions ($k$ = 0, 1, 2,,,, and $2_{j\text{-}1}$-1, the length of each of $j$ time-regions is $m/2_j$). Each of $2_j$ sub time-regions was multiplied with each of $2_j$ independent log-normal noises with length $m/2_j$ generated by $\exp\left[\omega^{(j,k)}\right]$. Absolute values of the generated multifractal time series are shown in Figure 4B.

**Movie preparation after recording**

A series of images (964 × 726 pixels, 8-bit depth) was acquired at a 50-ms interval and saved in a non-compressed tiff format by Micro-Manager camera control software (https://micro-manager.org). One thousand sets of 10,000 frames were recorded due to the limitations of Micro-Manager. The gap time intervals between each 10,000-frame recording was 0.5 s (10 frames), which was compensated for in the activity time series as described below. Each 10,000-frame recordings was processed automatically for removal of the background intensity gradient (Fig. S3A) and then compressed by an H265 codec (ISO/IEC 23008-2) in FFmpeg (https://www.ffmpeg.org), and saved in a 4 terabyte-hard disk during the recording. Compression parameters selected to balance compression efficiency and image quality were as follows: command line -c:v libx265 -threads 4 -x265-params qp=23:min-keyint=20 -preset:v veryfast. Artifactual activity due to compression was compensated for, as described below. The compression reduced 7 terabytes into 10–20 gigabytes (<0.2% non-compressed tiff data).

The brightness gradient at the radial axis in the image (illumination background) was removed by Gaussian filtering in Python scikit-image with a standard deviation of 7 pixels and 100 Gaussian-filtered images in the initial set of 10,000-frame recordings were averaged (Fig. S3B). Then, the averaged image was fit with two-dimensional fourth-degree polynomial function with image center as the origin point. All recorded images were subtracted with two-dimensional functions by fitting to obtain background-free images (Fig. S3C).

**Extraction of swimming activity time series and removal of artifactual swimming activity**

*C. elegans* movements were measured by counting the number of pixels that showed a significant intensity change within a 50-ms time interval due to *C. elegans* motions (Fig. 2A and B). To measure intensity changes in individual culture chambers, we manually made a circular mask to define area of a culture chamber in Inkscape vector graphics software (https://inkscape.org). According to the mask coordinates, an image difference defined by image[t +1] – image[t] in each of chamber was calculated to obtain swimming activity time series.

Occasionally, an air bubble moved through and was retained in the buffer flow, which caused activity artefacts in the pixel counting method. Moving bubbles affected local time series in chambers one by one as the bubble movement followed the buffer flow (Fig. S3D), whereas retained bubbles caused a continuous effect due to ruffling of the bubble edge and may affect chamber buffer exchange (Fig. S3F). In addition, accidental physical shock to the recording system e.g. from daily lab activities can cause surface ripples in the M9 buffer in the microfluidic device, which can also mimic artificial swimming activity (Fig. S3G and H). Moving bubbles and ripples were detected automatically based on a sudden reduction in mean pixel intensity within chambers (Fig. S3E) or in the average entropy of image differences among all chambers (Fig. S3H), respectively. To estimate image entropy, we computed $-\sum_{i=1}^{100} p_i log_2 p_i$, where $p_i$ for the $i$th bin was obtained from a normalized 100-bin histogram. The normalized histogram was computed within the region of each chamber in a 0–50 range of pixel intensity values after replacing negative image difference values with 0. To compensate for artifactual swimming activity, artificial animal activities were replaced with activities from adjacent recording frames (Fig. S3I). For replacing, a time region with artifactual swimming activities was divided into early and late halves, and was filled with the maximal activity value from among the activity data recorded within four time frames adjacent to each half (Fig. S3I). The maximal activity value was appropriate for estimates in the active or inactive state. The same replacing method was applied to the gaps between 10,000-frames sets due to Micro-Manager limitations and loss of movie files. For retained bubbles, we took the average minimum intensity of the image at each of 10,000 frames, which detect slow movement of bubbles, which appear as dark objects (Fig. S3F). We eliminated the whole activity time series of animals whose chambers had long-term retained bubbles. Finally, we obtained corrected-swimming activity time series of animals with 10,010,000 time points (= 10,000-frames × 1,000 sets of movie files + 1,000 × 10 gap frames).

Artifactual swimming activity caused by the intrinsic nature of the image compression codec was compensated for after obtaining activity time series from movies. H265 compressed a series of non-compressed tiff images into a movie file composed of an Intra-coded frame (I-frame), Predicted Frame (P-frame), Bi-directional Predicted Frame (B-frame) series [12]. H265 compresses and reconstructs the series of images with inter frame prediction; I-frame contains all the image information, P-frame is reconstructed by forward prediction from I-frame, and B-frame is reconstructed by forward and backward prediction from I- and P-frames. In our setting, the B-frame was responsible for a set of continuous four frames (BBBB, 0.2 s), and the P-frame was inserted after the continuous B-frame set. The I-frame was inserted after 50 BBBBP sequence repeats. To test whether temporal changes in *C. elegans* posture were properly reconstructed, we compared activity time series from H265-compressed images with that obtained from non-compressed tiff images and found that the Image[t +1] – image[t] differences were globally indistinguishable (Suppl. movie S5), though swimming duration tended to be truncated within a subsecond time scale (Fig. S6A). These slight truncations were attributed to a B-frame reconstruction insufficiency because an insufficient reconstruction of a difference in B-frames yields a zero or near-zero activity value. Quantitative comparisons of activity time series confirmed that H265-derived activity time series were most similar to tiff-derived series when the former were subjected to 3- and 5-frame smoothing in two different distance measures (Fig. S6B). The high similarity after 3-frame smoothing in H265-derived activity time series was likely caused by removal of Poisson noise because 3-frame smoothing reduces the effect of noise without a time correlation with adjacent frames. We found that 5-frame smoothing in H265-derived activity time series compensated well for slight truncations caused by B-frame reconstruction insufficiency. Based on the intrinsic characteristics of B-frame and experimental data analysis, we concluded that the strict limit for detecting *C. elegans* swimming motion in our setting was 5 frames (0.25 s). Swimming bouts longer than 0.25 s were thus deemed reliable. In addition, we convoluted swimming activity with a 5-frame rectangular function to compensate for H265-related slight truncations.

### References


1 Iannaccone, P. M. & Khokha, M. *Fractal Geometry in Biological Systems: An Analytical Approach*. (CRC Press, 1996).
2 Mandelbrot, B. B. *The fractal geometry of nature*. Updated and augm. edn, (W.H. Freeman, 1983).
3 Bunde, A. & Havlin, S. *Fractals in Science*. (Springer-Verlag, 1994).
4 Gneiting, T. & Schlather, M. Stochastic models that separate fractal dimension and the Hurst effect. *Siam Rev* **46**, 269-282, doi:10.1137/S0036144501394387 (2004).
5 Kiyono, K. Establishing a direct connection between detrended fluctuation analysis and Fourier analysis. *Phys Rev E Stat Nonlin Soft Matter Phys* **92**, 042925, doi:10.1103/PhysRevE.92.042925 (2015).
6 Kantelhardt, J. W. *et al.* Multifractal detrended fluctuation analysis of nonstationary time series. *Physica A* **316**, 87-114 (2002).
7 Ihlen, E. A. Introduction to multifractal detrended fluctuation analysis in matlab. *Front Physiol* **3**, 141, doi:10.3389/fphys.2012.00141 (2012).
8 Frisch, U. *Turbulence*. (Cambridge University Press, 1995).
9 Beran, J., Whitcher, B. & Maechler, M. longmemo; Statistics for Long-Memory Processes (Book Jan Beran), and Related Functionality, R package Version 1.1-1. (2018).
10 Ihaka, R. & Gentleman, R. R: A Language for Data Analysis and Graphics. *Journal of Computational and Graphical Statistics* **5**, 299-314, doi:10.2307/1390807 (1996).
11 Davies, R. B. & Harte, D. S. Tests for Hurst Effect. *Biometrika* **74**, 95-101, doi:DOI 10.1093/biomet/74.1.95 (1987).
12 Bing, B. *Next-Generation Video Coding and Streaming*.
