## Supplementary Figure legend for "*C. elegans* episodic swimming is driven by multifractal kinetics"

### Figure S1 self-similar and scale-free property in Koch curves and trajectory of a Brownian particle

(A) Koch curves drawn from line segments with length 1. Each elemental wedge shape is drawn at the line-segment center and smaller wedges are drawn in each remaining shorter segment. This process is repeated infinitely. Koch curve lengths are changed with spatial resolution for measurement $((4/(2+\sqrt{3}))^n$, $(4/3)^n$, $2^n$, respectively; $n$, arbitrary resolution unit). Fractal dimensions of each Koch curves are indicated below. (B) Numerically-generated Brownian particle trajectory was magnified sequentially ten times in a lower time series.

### Figure S2 Buffer exchange in WormFlo

(A) Fluorescence image of WormFlo with saturated 10 μM FITC solution in culture. (B) Fluorescence image of the optical field with 10 μM FITC solution without WormFlo used for a reference to estimate a background intensity gradient in the radial axis due to optical limitations (e.g. biased illumination). Despite having a uniform concentration, 10 μM FITC solution exhibited a background intensity gradient. (C and D) Mean fluorescence intensities in the chambers surrounded with the yellow circle decayed as FITC-free waster was supplied, as shown in linear (C) and semi-log (D) plots. Fluorescence intensity within a chamber was measured by subtracting background intensity gradient (B) and the fluorescence intensity in the flow path under the chamber (laterally adjacent area to the chamber). The decay kinetics have at least two rate constants: a fast rate due to soluble FITC (half-time, ~250 s), and a slow rate likely due to photobleaching of FITC absorbed into PDMS (D).

### Figure S3 Temperature and light control in the recording system

(A and B) For temperature maintenance, the WormFlo device was submerged in M9 buffer in a 15-cm-diameter glass dish, which was covered by aluminum box (A) attached to an underlying aluminum board with a 1-cm-diameter water flow path (B). Temperature controlled water perfuses continuously into the water path in the aluminum board. (C) The temperatures of the laboratory room (yellow), light shield box (green), water in the glass dish (red), and temperature-controlled water bath (blue) were tracked by a temperature logger (TC-08, Pico technology, UK). Temperatures in the glass dish containing the WormFlo were maintained within 0.5 ˚C. (D) Spectra of light at macroscope stage with (orange line) or without (black line) a light-filtering orange acryl board, in the presence (orange and black lines) or absence of illumination (gray line). Spectrum shapes show filtering out of light with wavelengths < 500 nm by the orange acryl board (shift in orange vs. black lines), and laboratory room fluorescent light (black line, lower panel) did not affect light intensity on WormFlo. Light spectrum with an orange acryl board (orange line) represents the light used for observation. (E) Mean light intensity at WormFlo culture chambers over 6 d (standard deviation <0.1% of the mean).

### Figure S4 Image processing to remove illumination background and detect air bubbles in the flow channel and accidental physical shocks to the recording system

Non-compressed tiff images (8-bit depth) had a radial-axis intensity gradient due to illumination light properties (A). The intensity gradient was removed by an image mask (B) made from recorded images by Gaussian filtering-based method (Supplemental information) to obtain background-removed images (C). (D) Air bubble in the flow path (red arrow). (E) Moving air bubbles were detected by a sudden drop in average pixel intensity in a chamber (black arrow) caused by their dark periphery. (F) Non-moving "staying" bubbles were detected by averaging intensities under a threshold to detect the staying bubble edges (E) among $10^3$ images obtained in every $10^4$-frame interval over the entire recording of $10^7$ frames. (G and H) M9 buffer surface ripples due to accidental physical shocks to the recording system were detected by a sudden rise in mean image entropy estimates among 108 culture chambers [red (G) and black (E) arrows]. Image entropy estimates were computed as $-\sum_{i=1}^{100} p_i log_2 p_i$, where $p_i$ for $i^{th}$ bin was obtained from a normalized 100-bin histogram of pixel intensities in image difference with the positive values from 0 to 50 in each chamber (see Fig. 2A), which is represented in a probability density function with the bin size of 0.5 (50 intensity intervals/100 bins). Due to the non-unity bin size, image entropy estimates can have a negative value. (I) Raw swimming activity in a frame in which actual swimming activity (red) could not be recorded due to air bubbles or ripples was compensated with activities recorded in adjacent time frames (blue), as indicated by the green bracket.

### Figure S5 Swimming activity of wild-type animals cultured with an energy source or *egl-4* mutants cultured without energy source

(A) Sustained swimming activity observed in a representative sample (5 of 108) of wild-type animals cultured with glucose and cholesterol in M9 buffer. Sustained swimming activity exceeding the High activity class threshold without an energy source was observed in 35/108 animals. The High:Middle:Low ratios of swimming activity classes, in %(N), were 73.1% (79):16.6% (18):10.2% (11) with an energy source and 30.7% (20):10.8% (23):33.9 % (22) in M9 buffer alone. The portion of High activity animals was more than doubled with an energy source versus that without ($p = 1.17 \times 10^{-7}$, chi-squared test for independence). (B) Swimming activity of a representative *egl-4* mutant at various time scales; under the full recording of $10^7$ timepoints (top) are shown several-day, several-hour, about-an-hour, several-minute, about-a-minute, and several-second time scales. Colored areas above indicate Pre-starved (red) and Starved (blue) time regimes. Magnifications of the first tenth of each upper panel are shown below.

**Figure S6 Estimation of artifactual effect on activity time series due to movie compression**

Pixel areas where animals moved were detected as image differences of non-compressed tiff images (red) and H265-compressed images (green) were superimposed on bright-field images (grey). The global trends in swimming activity strength in non-compressed tiff images (red) were represented in the H265-compressed images (upper panel). As a minor difference, continuous swimming periods in H265-compressed images tended to be locally truncated (red arrows in lower panel). Active state (high blue-line value) and inactive state (low blue-line value) were determined by a threshold at 12 pixels/frame. (B) Swimming activity time series of 108 animals; two distance measures (correlation coefficient and Euclidian distance) were compared between10,000 frames of non-compressed tiff images versus H265-compressed images. Correlation coefficient and Euclidian distance values obtained from tiff-derived activity time series after rectangular convolution among 3-frames were compared with H265-derived activity time series after rectangular convolutions among various frames (window size in x-axis). Subsequently, rectangular convolutions among 5-frames showed a high correlation, implying that the temporal truncation of activity time series due to image compression was compensated with rectangular convolutions among 5-frames by smoothing effect in convolution. (C) Activity time series compensated with 5-frame rectangular convolutions (compressed+smoothing). Each activity time series was obtained by a pixel counting method where active pixels were identified based on image differences between original tiff images and H265-compressed images (original vs. compressed). Each series was convoluted with 5-frame rectangular convolutions (original+smoothing and compressed+smoothing). Post-compression and smoothing activity time series were analyzed in this study.

**Movie S1 Movie for wild-type animals cultured with M9 buffer in WormFlo**

Swimming activity was recorded in wild-type animals cultured in a WormFlo without an energy source. Pixel areas where animals moved were detected by image differences of H265-compressed images (green) and were indicated on bright field of images (grey) (upper panel). Swimming activity of animal ID 43 indicated by a red frame was quantified from a series of H265-compressed images (green) by a pixel counting method (lower panel). The swimming activity (vertical dashed line) at each time point indicates the image difference shown in the upper panel. Posing occurs around time points (in s) 2, 3, 14, 16, 21, 26, and 28.5.

**Movie S2 $10^4$ times fast-forward movie of wild-type animals cultured with M9 buffer in WormFlo**

Swimming activity was recorded in wild-type animals cultured in a WormFlo without an energy source and played at a 10000× fast-forward rate.

**Movie S3 $10^4$ times fast-forward movie of wild-type animals cultured with M9 buffer in 96-well plate wells**

Swimming activity was recorded in wild-type animals cultured in wells in a 96-well plate without an energy source and played at a 10,000× fast-forward rate.

**Movie S4 $10^4$ times fast-forward movie of wild-type animals cultured with an energy source in WormFlo**

Swimming activity was recorded in wild-type animals cultured with an energy source in a WormFlo and played at a 10,000× fast-forward rate.

### Movie S5 Comparison of swimming activity obtained from a series of H265-compressed images and non-compressed tiff images

Pixel areas where *egl-4(n479)* mutant animals cultured in a WormFlo without an energy source move in a movie frame were detected as image differences of H265-compressed images (green) versus original non-compressed tiff images (red) (upper panel). Swimming activity of animal ID 9 was quantified from a series of H265-compressed images (green) and original non-compressed tiff images (red) by a pixel counting method (lower panel). Swimming activity at each time point (vertical dashed line) corresponds to the image difference shown in the upper panel. Active versus inactive states were distinguished at a threshold of 12 pixels/frame (horizontal line).
