## Supplementary figures and images for "*C. elegans* episodic swimming is driven by multifractal kinetics"

### Figure S1

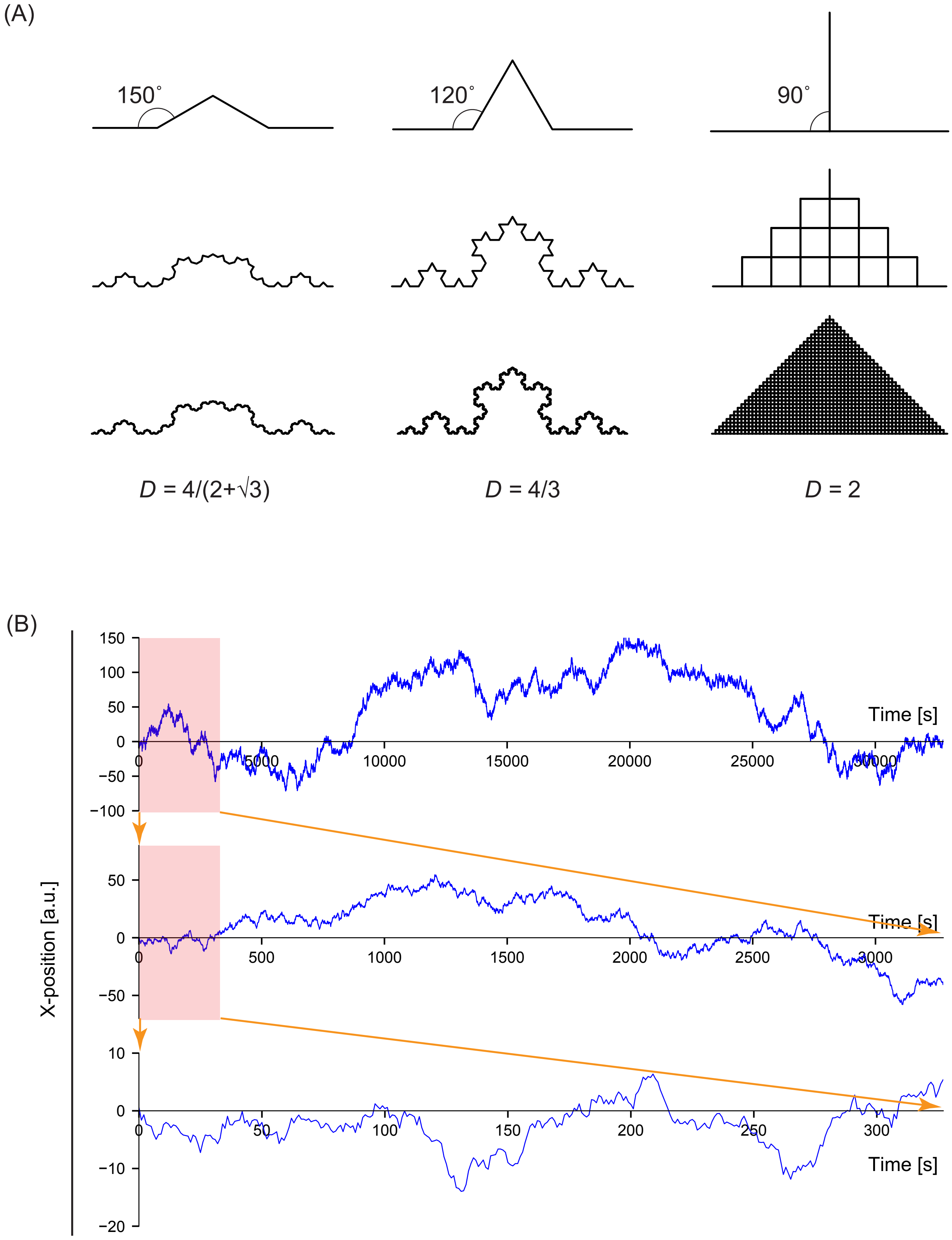


Figure S1 Ikeda et al

### Figure S2

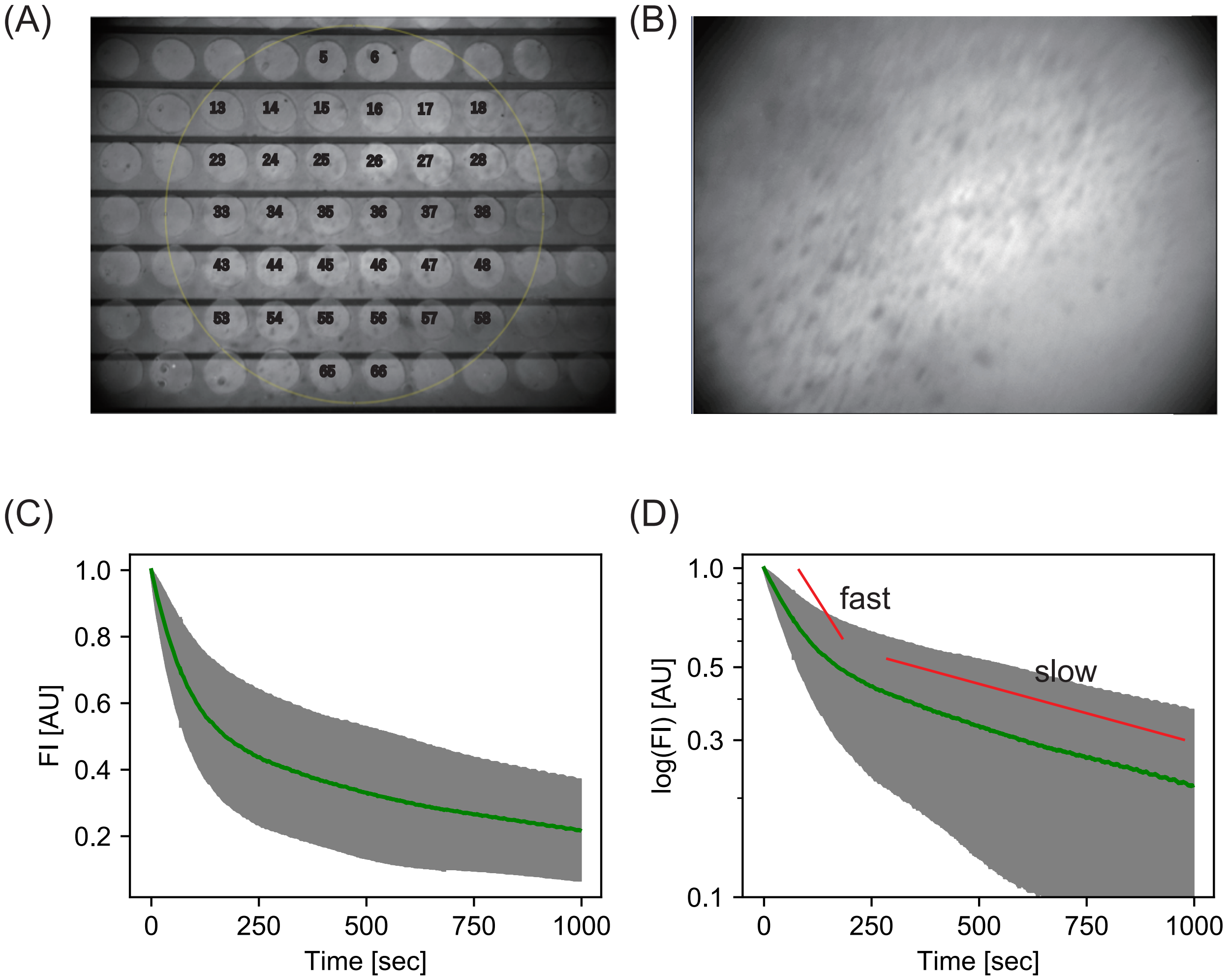


Figure S2 _IKeda et al

### Figure S3

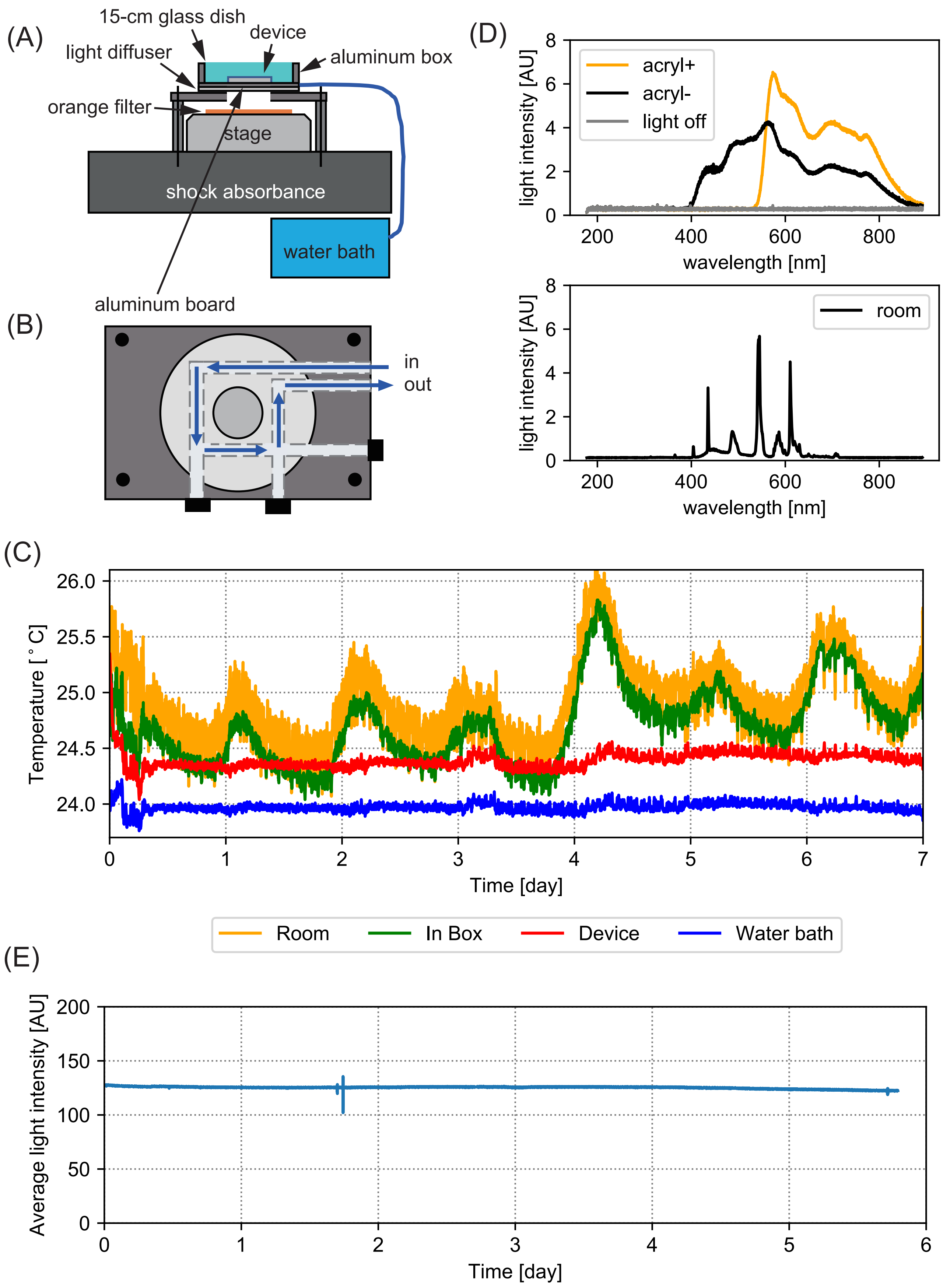


Figure S3_Ikeda et al

### Figure S4

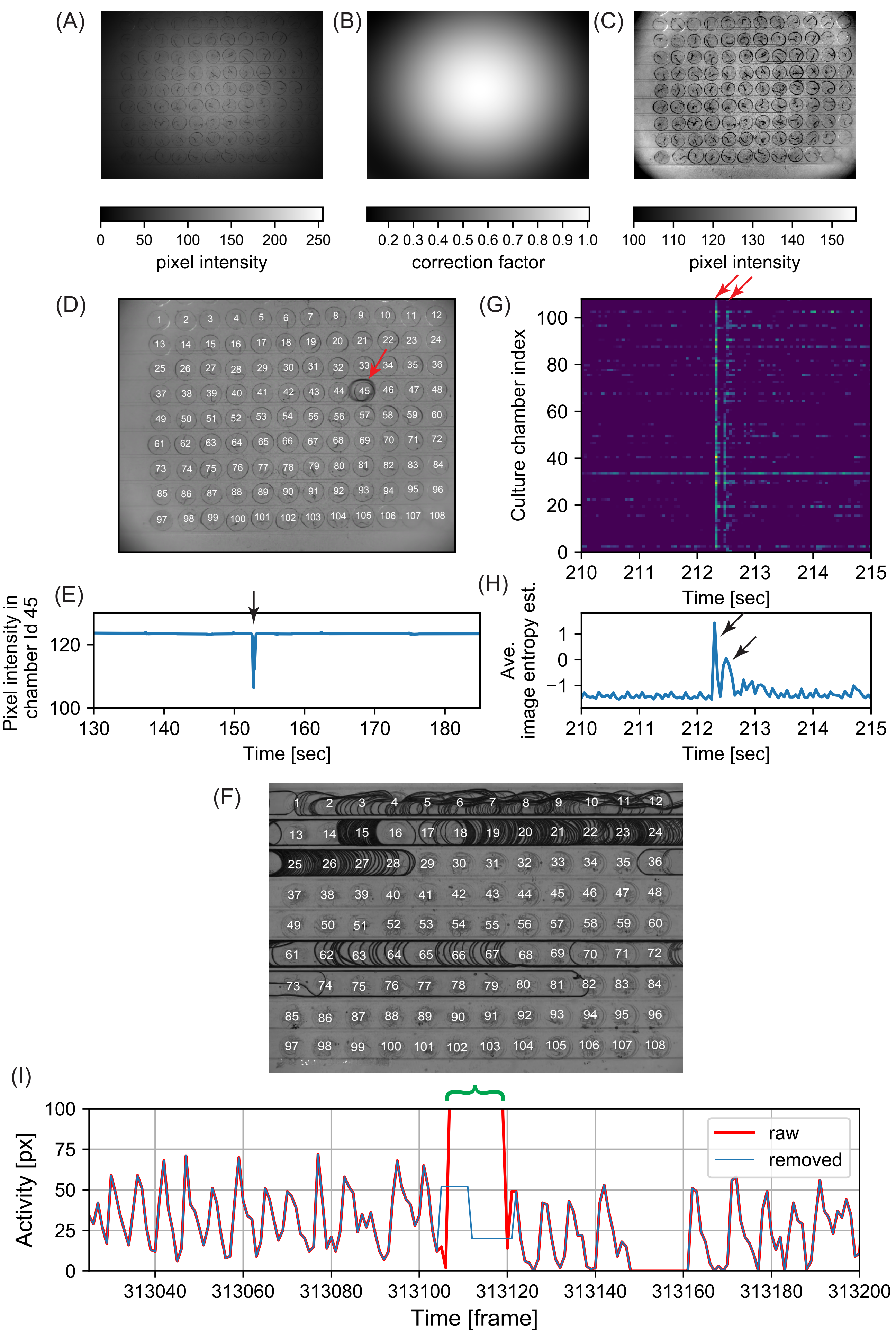


Figure S4_Ikeda et al

### Figure S5

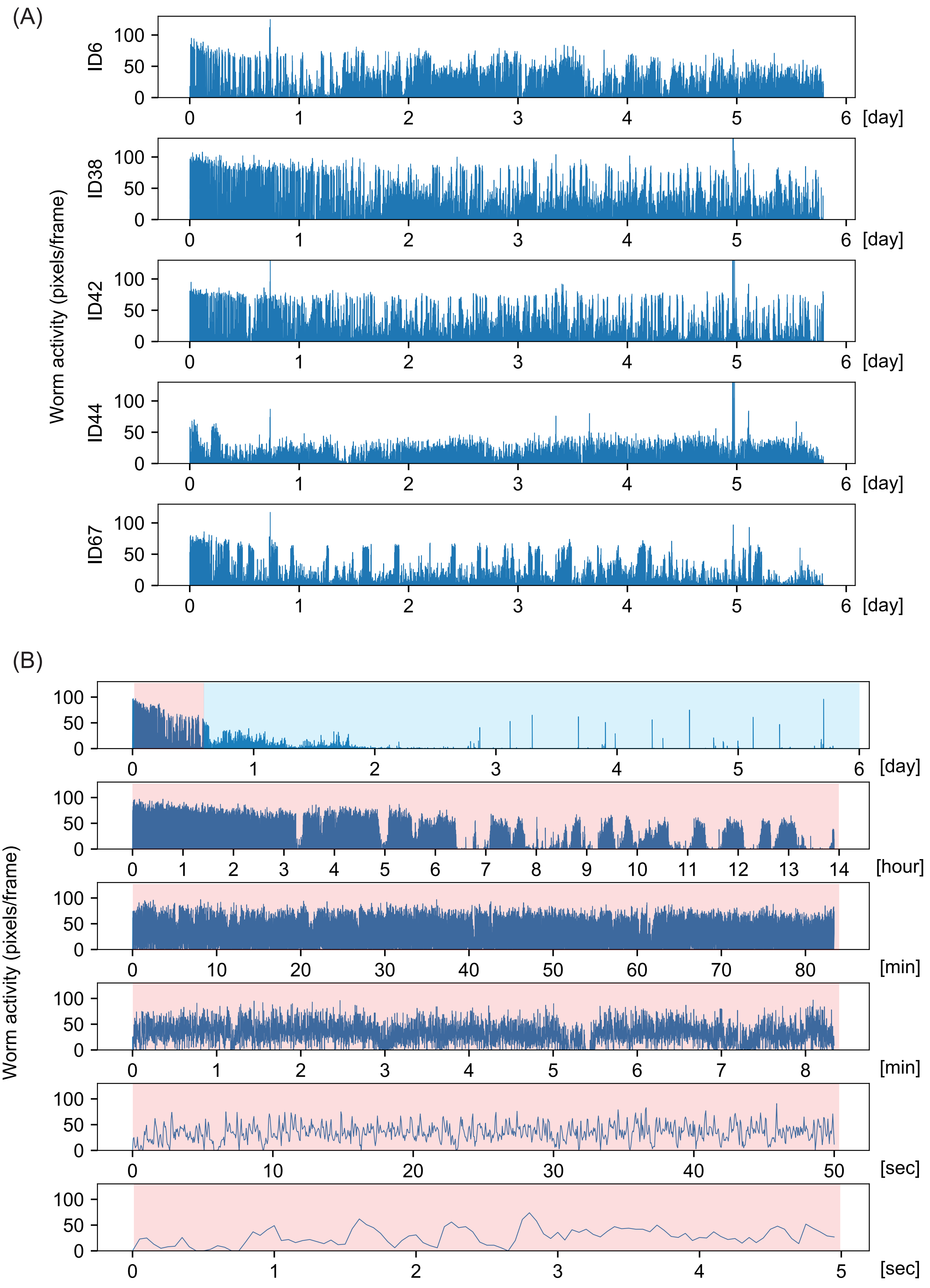


Figure S5_Ikeda et al

### Figure S6

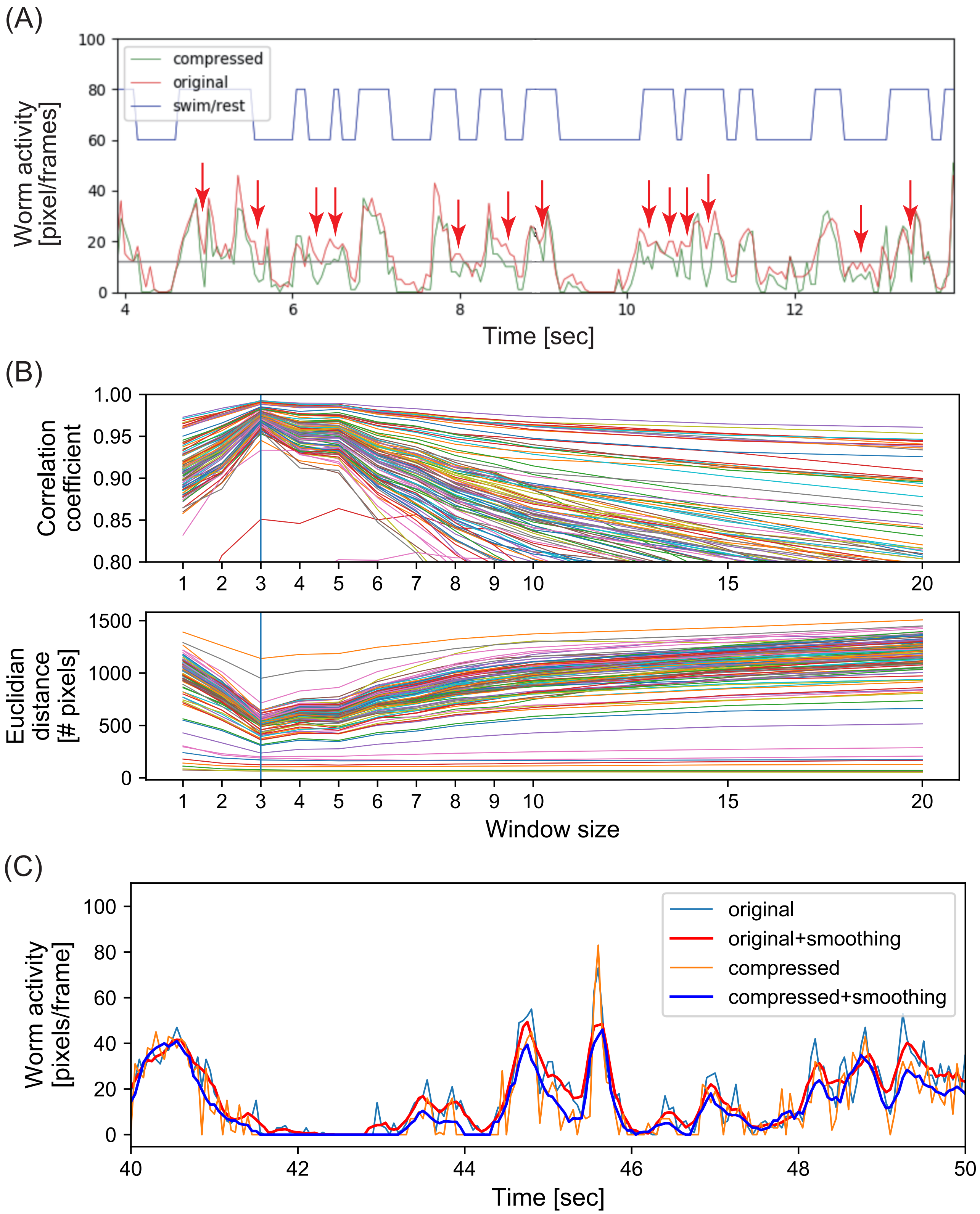


Figure S6 Ikeda et al
