## Supplementary material for "*C. elegans* episodic swimming is driven by multifractal kinetics": Tables

**Table 1** Summary of fit parameters for power law distributions in Figure 3D–G and t-test p-values for the slope of power law distribution fit parameters.

**fit parameters**

| | | Active | | Inactive | |
|---|---|---|---|---|---|
| | | *a* | *b* | *a* | *b* |
| High | Pre-starved | 1.89 ± 0.17 | 0.81 ± 0.07 | 1.62 ± 0.13 | 1.00 ± 0.13 |
| | Starved | 1.76 ± 0.57 | 1.52 ± 0.21 | 1.20 ± 0.23 | 0.90 ± 0.18 |
| Middle | Pre-starved | 1.81 ± 0.20 | 0.77 ± 0.12 | 1.67 ± 0.20 | 1.04 ± 0.20 |
| | Starved | 2.15 ± 0.74 | 1.30 ± 0.34 | 1.34 ± 0.24 | 0.92 ± 0.12 |
| Low | Pre-starved | 1.87 ± 0.28 | 0.80 ± 0.17 | 1.65 ± 0.21 | 1.06 ± 0.22 |
| | Starved | 2.06 ± 0.76 | 1.28 ± 0.34 | 1.30 ± 0.32 | 0.93 ± 0.20 |
| *egl-4(n479)* | Pre-starved | 2.02 ± 0.43 | 0.65 ± 0.16 | 1.96 ± 0.13 | 1.04 ± 0.19 |
| | Starved | - | - | - | - |

**p-values**

| | Active vs Inactive | | Pre-starved vs Starved |
|---|---|---|---|
| **High activity class** | | **High activity class** | |
| Pre-starved | $7.9\times10^{-6}$* | Active | $5.70\times10^{-1}$ |
| Starved | $2.6\times10^{-2}$* | Inactive | $5.07\times10^{-4}$* |
| **Middle activity class** | | **Middle activity class** | |
| Pre-starved | $3.1\times10^{-3}$* | Active | $3.03\times10^{-2}$* |
| Starved | $1.2\times10^{-5}$* | Inactive | $2.61\times10^{-7}$* |
| **Low activity class** | | **Low activity class** | |
| Pre-starved | $4.9\times10^{-6}$* | Active | $1.50\times10^{-1}$ |
| Starved | $8.0\times10^{-7}$* | Inactive | $1.22\times10^{-7}$* |
| ***egl-4(n479)*** | | ***egl-4(n479)*** | |
| Pre-starved | $4.2 \times 10^{-1}$ | Active | - |
| Starved | - | Inactive | - |

| | *egl-4(n479) vs* High activity class |
|---|---|
| Active | $1.09 \times10^{-1}$ |
| Inactive | $7.79\times10^{-11}$* |

(Upper table) Residence time distributions of animals' active and inactive states in Pre-starved and Starved time regimes (defined as in Fig. 2) on a log-log scale fit with y = $-ax-b$. Mean $a$ and $b$ values are shown with the standard deviations. (Lower table) $p$-values from t-tests of differences in slope $a$ from upper table in indicated comparisons; $*p < 0.05$.

**Table 2** Summary of $H_{peak}$ and *width* values of the multifractal spectra in Figure 5A–D and associated t-test p-values.

**$H_{peak}$ and *width***

| | | Active | | Inactive | |
|---|---|---|---|---|---|
| | | $H_{peak}$ | *width* | $H_{peak}$ | *width* |
| High | Pre-starved | 0.89 ± 0.05 | 0.67 ± 0.11 | 1.39 ± 0.14 | 1.49 ± 0.19 |
| | Starved | 0.83 ± 0.09 | 0.50 ± 0.14 | 1.34 ± 0.39 | 1.88 ± 0.29 |
| Middle | Pre-starved | 0.88 ± 0.09 | 0.77 ± 0.21 | 1.30 ± 0.31 | 1.51 ± 0.20 |
| | Starved | 0.88 ± 0.13 | 0.62 ± 0.19 | 1.34 ± 0.60 | 2.00 ± 0.35 |
| Low | Pre-starved | 0.80 ± 0.10 | 0.65 ± 0.18 | 1.33 ± 0.36 | 1.65 ± 0.41 |
| | Starved | 0.81 ± 0.18 | 0.61 ± 0.32 | 1.59 ± 0.33 | 2.25 ± 0.81 |
| *egl-4(n479)* | Pre-starved | 0.88 ± 0.07 | 0.53 ± 0.20 | 1.13 ± 0.20 | 1.28 ± 0.25 |
| | Starved | - | - | - | - |

**p-values**

| | Active vs Inactive | | | Pre-starved vs Starved | |
|---|---|---|---|---|---|
| | $H_{peak}$ | *width* | | $H_{peak}$ | *width* |
| **High activity class** | | | **High activity class** | | |
| Pre-starved | $9.52 \times 10^{-13}$* | $7.51 \times 10^{-16}$* | Active | $1.36\times10^{-1}$ | $1.06\times10^{-2}$* |
| Starved | $6.50 \times 10^{-3}$* | $8.16\times10^{-8}$* | Inactive | $7.07\times10^{-1}$ | $5.33\times10^{-3}$* |
| **Middle activity class** | | | **Middle activity class** | | |
| Pre-starved | $1.00 \times 10^{-6}$* | $2.38\times10^{-16}$* | Active | $8.33\times10^{-1}$ | $2.33\times10^{-2}$* |
| Starved | $6.70 \times 10^{-3}$* | $3.61\times10^{-13}$* | Inactive | $8.06\times10^{-1}$ | $3.70 \times10^{-5}$* |
| **Low activity class** | | | **Low activity class** | | |
| Pre-starved | $1.00 \times 10^{-5}$* | $3.18\times 10^{-9}$* | Active | $8.74\times10^{-1}$ | $7.76\times10^{-1}$ |
| Starved | $2.00 \times 10^{-6}$* | $1.91\times10^{-5}$* | Inactive | $5.92\times10^{-2}$ | $3.27\times10^{-2}$* |
| ***egl-4(n479)*** | | | ***egl-4(n479)*** | | |
| Pre-starved | $4.45\times10^{-9}$* | $1.98\times10^{-23}$* | Active | - | - |
| Starved | - | - | Inactive | - | - |

| | *egl-4(n479)* vs wild-type High | |
|---|---|---|
| | $H_{peak}$ | *width* |
| Active | $5.42\times10^{-1}$ | $1.15\times10^{-3}$* |
| Inactive | $1.00\times10^{-6}$* | $1.16\times10^{-3}$* |

(Upper table) Hurst exponent for the entire time series ($H_{peak}$) and widths of multifractal spectrum (distance of local Hurst exponent at -10 < $q$ < 10) obtained from individual animals in the indicated activity class or mutant animals in Pre-starved and Starved time regimes. (Lower table) ) p-values from t-tests of differences in $H_{peak}$ and *width* from upper table in indicated comparisons; *p < 0.05.
